# The Spatial Landscape of Extracellular Matrix Gene Expression in Healthy and Type 2 Diabetic Human Pancreas

**DOI:** 10.64898/2026.06.30.733275

**Authors:** Luisa Kimberly Meneses, Honesty Jung Kim, Gregory L. Szot, Julie B. Sneddon, Zev J. Gartner

## Abstract

The unique peri-islet and double-layered vascular basement membrane (BM) of the human pancreatic islet are critical regulators of beta cell survival and function. While animal models imply that endothelial cells (ECs) are the exclusive source of islet BM, the precise cellular origins and spatial organization of the human islet matrisome remain poorly defined due to overlap in genes that mark non-epithelial cell populations and loss of spatial context during single-cell dissociation. In this study, we combine computational integration of whole-pancreas single-cell transcriptomes using CONCORD with high-resolution MERFISH spatial genomics to map the extracellular matrix (ECM) landscape across 251,477 spatially resolved cells from seven non-diabetic and five type 2 diabetic human donors. Contrary to an endothelial-centric paradigm, our data support a cooperative division of labor in the provision of BM, where pericytes represent the dominant transcriptional source of structural BM collagens (*COL4A1*, *COL4A2*) and ECs selectively express complementary matrix factors (*HSPG2*, *LAMA5*). Spatial neighborhood analysis further resolves a specialized population of islet-associated fibroblasts enriched at the islet boundary that are characterized by expression of peri-islet laminin genes. In type 2 diabetes, this homeostatic perivascular niche changes composition, marked by a significant increase in the islet fibroblast-to-pericyte ratio. Concurrently, islet pericytes undergo pro-fibrotic reprogramming characterized by the loss of canonical identity markers (*PDGFRB*), altered expression of ECM genes including *COL1A2* and *COL18A1*, and upregulation of contractile machinery (*MYL9*). In the non-diabetic pancreas, pericytes constitute the principal vascular BM-expressing population within islets, whereas type 2 diabetes is associated with coordinated, compartment-specific remodeling of vascular-supportive stromal populations.

**Research in Context:** *What is already known about this subject?:*

- Extracellular matrix (ECM), and in particular basement membrane (BM), are essential structural and signaling components of the pancreatic islet microenvironment that contribute to beta cell function and survival.
- Islet capillaries are closely associated with endocrine cells and are surrounded by specialized BMs; however, the cellular sources of these BM components in the adult human pancreas remain incompletely defined.
- Type 2 diabetes is associated with islet fibrosis and vascular dysfunction, but cell type-specific alterations in ECM-producing populations have not been comprehensively characterized *in situ*.

*What is the key question?:*

- Which cell populations produce the components of ECM, including BM, within the adult human islet, and how are these populations altered in type 2 diabetes?

*What are the new findings?:*

- Spatial transcriptomics identifies pericytes as the predominant vascular-associated source of ECM, including BM, gene expression in human islets, whereas endothelial cells exhibit complementary but more limited matrix-producing programs.
- Spatial transcriptomics identifies an islet-associated fibroblast population enriched for fibrillar collagen and BM-associated genes that localizes preferentially to the islet surface niche.
- Type 2 diabetes is associated with remodeling of perivascular ECM programs, including reduced expression of vascular basement membrane genes, a shift from a pericyte to smooth muscle-like identity, and increased expression of matrix-remodeling and fibrosis-associated genes.

*How might this impact clinical practice in the foreseeable future?:*

- Defining the cellular sources and disease-associated remodeling of the human islet ECM may inform the development of therapies aimed at preserving or restoring the islet microenvironment in type 2 diabetes.
- Including key subtypes of islet-associated ECM-producing cells may be important in improving current protocols to generate replacement islets from human pluripotent stem cells for cell replacement therapy for diabetes.

## Introduction

The islets of Langerhans are clusters of endocrine cells, distributed across the exocrine pancreas, that play a central role in glucose homeostasis. Islets are heterogeneous in size (20–500 µm in diameter) and are surrounded by a specialized extracellular matrix (ECM) structure known as the peri-islet basement membrane (BM), which separates the endocrine and exocrine compartments[1–3]. A second source of BM surrounds the intra-islet vasculature, which in humans is uniquely organized as a double layer[4]. Unlike the broader interstitial ECM, which includes fibrillar collagens, fibronectins, and proteoglycans, BM is a thin ECM sheet composed primarily of laminins, collagen IV, nidogen, and perlecan that sits in direct contact with the epithelial and endocrine surfaces[5–7]. It is through this direct interface that BM conveys adhesion, polarity, and survival signals to the cells it surrounds[5, 8]. In the islet, these two regionally distinct BM sources provide critical structural and instructive cues to the endocrine cells, additionally influencing insulin secretion, gene expression, and spatial organization both *in vivo* and *in vitro* [9–17]. These regions further differ in their ECM protein composition, suggesting functional specialization and potentially distinct cellular origins that remain poorly understood [2–4, 18]. Defining which cells produce the islet BM and which laminin and collagen isoforms they express is therefore fundamental to understanding how the beta cell niche is built and maintained, with implications for modeling disease, understanding the progression of diabetes, and engineering ECM scaffolds for cell replacement therapy [1, 4, 19].

Studies in the mouse have established that endothelial-deficient islets lack intra-islet BM, suggesting that, unlike many epithelial cell types, endocrine cells do not synthesize their own BM proteins[13]. Instead, these studies have suggested that beta cells signal to endothelial cells (ECs) to promote vascularization and BM deposition during islet development, leading to the prevailing view that ECs are the main source of islet BM[13]. However, emerging evidence challenges this notion[10, 17]. Perivascular cells, particularly pericytes, are integral components of the vascular niche and may contribute to vascular BM assembly in addition to ECs. Published human vascular transcriptomic datasets and experimental vascular models have demonstrated that pericytes express BM-associated genes and participate in vascular matrix deposition[20, 21]. Mouse studies further indicate that pancreatic pericytes contribute to islet BM formation and regulate beta cell function by modulating blood flow and BMP4 signaling[22–24]. RNA sequencing has further revealed that mouse islet pericytes express a substantial number of ECM-encoding genes, sometimes at levels comparable to or exceeding those of other islet cells[17]. However, whether these findings extend to humans remains unknown.

Given the unique organization of the human islet compared to mouse, it is critical to understand the specific roles of the epithelial, endothelial, and stromal cells in constructing the double-layered vascular BM and peri-islet BM. Filling this gap has been challenging because vascular cells exhibit organotypic transcriptional features, including the expression of ECM-encoding genes[25–27]. For instance, pancreatic ECs present in the endocrine compartment display distinct transcriptional profiles from those in the exocrine compartment [25, 28–31]. Islet-associated ECs upregulate *PLVAP*, a key regulator of fenestrated capillaries essential for efficient insulin release, highlighting the need to analyze endocrine and exocrine-associated vascular cells independently [25].

Beyond its important role in pancreatic development and function, BM remodeling is increasingly recognized as a feature of metabolic diseases such as type 2 diabetes[32, 33]. While most research in the field of islet biology in type 2 diabetes has focused on endocrine dysfunction, vascular alterations and ECM remodeling also play a role in disease pathology[32–38]. In type 2 diabetes, ECM modifications can impact endocrine cell adhesion, migration, and survival, ultimately disrupting islet function[32, 36, 37, 39–42]. Which cell types contribute to these changes in ECM remains unknown.

Identifying the sources of BM proteins in the non-diabetic and type 2 diabetic human pancreas requires studying vascular and perivascular cells within intact tissue. Since pericytes, ECs, and fibroblasts are distributed across endocrine, exocrine, and interstitial compartments, dissociation-based approaches like single-cell RNA-sequencing (scRNA-seq) have been unable to distinguish between different tissue regions alone. One solution to this problem has been isolating endocrine and exocrine tissues separately, then applying mechanical and enzymatic dissociation to prepare each tissue type for scRNA-seq. However, stripping cells of their spatial context, even temporarily, can alter the transcriptional states of ECM-expressing cells [18, 43–46]. Immunofluorescence (IF) offers the appropriate context and spatial resolution but is limited by the availability of specific ECM-targeting antibodies and significant marker overlap among vascular cells. Additionally, most ECM proteins are secreted and deposited adjacent to the cells in which they were synthesized. Therefore, IF cannot reliably trace matrix components back to their cellular source[39, 47].

To overcome these technical challenges, we combine careful integration of droplet scRNA-seq data together with MERFISH (Multiplexed Error-Robust Fluorescence *In Situ* Hybridization) technology on the Vizgen MERSCOPE platform, enabling highly multiplexed, single-cell resolution gene expression within intact tissues[48, 49]. Unlike conventional methods, MERFISH is highly sensitive and allows for the precise mapping of ECM gene expression while preserving the native architecture of the pancreas. This approach has enabled the simultaneous visualization of ECM gene expression patterns across thousands of cells and twelve human donors, validated against a gold-standard, unbiased, droplet-based single-cell transcriptional atlas of the human pancreas from non-diabetic and type 2 diabetic donors.

## Materials and Methods

### Human pancreas sample acquisition

De-identified blocks of formalin fixed, paraffin-embedded (FFPE) pancreatic tissue from 12 unremarkable, cancer-free adult cadaveric donors were purchased from ProteoGeneX (Inglewood, CA; RRID:SCR_013844) or collected from the University of California San Francisco Diabetes Research Center Islet Production Core Facility (RRID:SCR_015106). All donors’ families gave informed consent for the use of pancreatic tissue in research. Of these, 7 donors were classified to be non-diabetic, and 5 donors had a documented clinical diagnosis of type 2 diabetes. Disease status was determined from donor clinical metadata provided by the tissue source. One non-diabetic donor was obtained through UCSF Human Islet Core; the remaining donors were obtained from ProteoGeneX. Available donor demographic and clinical information are provided in Table 1 and detailed in Table S5.

**Table 1.** Human pancreatic donor cohort. Demographic and clinical information for non-diabetic and type 2 diabetic donors included in this study. Additional donor information is provided in Table S5.

| Sample ID | Cluster Label | Sex | Age | Cohort | Ethnicity | Type | Diagnosis | DV200 | BMI |
| --- | --- | --- | --- | --- | --- | --- | --- | --- | --- |
| 121771A(2) | 4 | M | 46 | ProteoGeneX | Caucasian | FFPE | Non-Diabetic | 34.65% | 29.4 |
| 121753A(1) | 5 | F | 41 | ProteoGeneX | Caucasian | FFPE | Non-Diabetic | 65.36% | 19.3 |
| 121467A(5) | 3 | M | 70 | ProteoGeneX | Caucasian | FFPE | Non-Diabetic | 56.55% | 29.3 |
| 121431A(4) | 6 | F | 73 | ProteoGeneX | Caucasian | FFPE | Non-Diabetic | 51.25% | 33.1 |
| 121320A(4) | 2 | M | 46 | ProteoGeneX | Caucasian | FFPE | Non-Diabetic | 65.76% | 26.9 |
| 121261A(3) | 8 | M | 69 | ProteoGeneX | Caucasian | FFPE | Type 2 Diabetes | 40.20% | 26.8 |
| 12997A(1) | 9 | F | 69 | ProteoGeneX | Caucasian | FFPE | Type 2 Diabetes | 62.93% | 31.2 |
| 12951A(1) | 10 | F | 74 | ProteoGeneX | Caucasian | FFPE | Type 2 Diabetes | 69.51% | 15.1 |
| 12893A(2) | 11 | M | 71 | ProteoGeneX | Caucasian | FFPE | Type 2 Diabetes | 61.74% | 24.7 |
| 12774A(1) | 12 | F | 58 | ProteoGeneX | Caucasian | FFPE | Type 2 Diabetes | 45.21% | 32.9 |
| 121903A(3) | 1 | M | 75 | ProteoGeneX | Caucasian | FFPE | Non-Diabetic | 57.08% | 30.2 |
| RHIP156 | 7 | F | 48 | UCSF | African American | FFPE | Non-Diabetic | 70.50% | 24.1 |

### Python analysis

All computational analyses were performed in Visual Studio Code (v.1.124; RRID:SCR_026031), Python (v3.8-v3.13) using Scanpy (v1.12; RRID:SCR_018139)[50], Squidpy (v1.6.2; RRID:SCR_026157)[51], scvi-tools (v0.6; RRID:SCR_026673)[52], DoubletDetection (v2.5.2), GSEApy (v1.2.1; RRID:SCR_025803)[53], SciPy (v1.17.1; RRID:SCR_008058)[54], Shapely (v2.1.2; RRID:SCR_027411), and CONCORD-sc (v1.11.4)[55] or Fiji/ImageJ (v2.16; RRID:SCR_003070)[56].

### scRNA-seq Quality Control and Filtering

scRNA-seq data (.h5ad) were downloaded from the PancDB (RRID:SCR_021860) website [57] and further filtered using Scanpy. Quality control was performed to exclude low-quality cells and technical artifacts. Cells with ‘unknown’ cell label, fewer than 100 detected gene counts, or greater than 20,000 detected gene counts were removed to eliminate empty droplets and potential doublets. Doublets were identified using DoubletDetection.

### Integration and Batch Alignments with CONCORD

To correct for donor- and dataset-specific batch effects, data were integrated using the CONCORD framework, incorporating donor/sample identity as a batch variable[55].

### Gene Set Scoring

Gene set enrichment was quantified at the single-cell level using sc.tl.score_genes. For whole-transcriptome analyses, the NABA Basement Membrane gene set obtained from MSigDB (RRID:SCR_016863) was used for BM gene set scoring. For spatial analyses, BM scores were calculated using a manually curated subset of BM-associated genes represented in the MERFISH gene panel (Figure 3A, Table S3).

**Figure 1:**
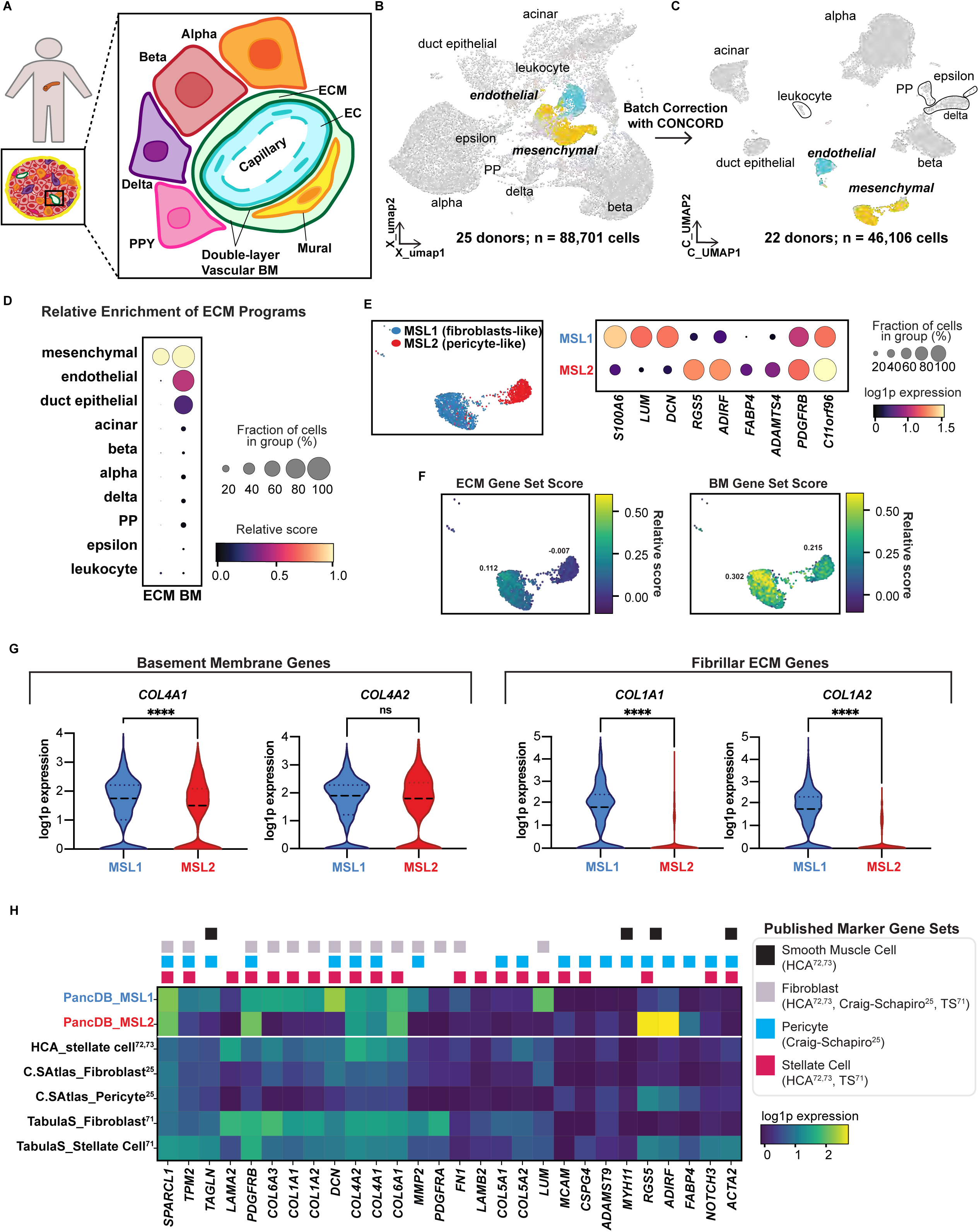
Mesenchymal cells are the dominant source of basement membrane transcripts in the healthy adult human pancreas. (A) Schematic of the pancreatic vascular niche depicting the specialized architecture of the islet microvasculature. Fenestrated endothelial cells (cyan) are encircled by perivascular mural cells (yellow) and sequestered within a distinct, double-layered vascular basement membrane (BM; green). Endocrine cells (red, beta cell; orange, alpha cell; purple, delta cell; pink, PPY cell) are in close physical contact with the islet vascular BM. (B) UMAP projection of PancDB single-cell atlas of the non-diabetic (ND) adult human pancreas prior to curation of data, showing 88,701 cells across 25 ND adult human donors visualized prior to batch alignment. Original published cell-type annotations are shown, with endocrine and exocrine populations in grey and candidate vascular niche populations, mesenchymal (yellow) and endothelial (cyan), highlighted for downstream analysis[70]. (C) Single-cell data from (B) after integration with CONCORD. UMAP projection of 46,106 cells across 22 ND adult human donors visualized following batch alignment, filtering of doublets and low-quality samples, and integration with CONCORD. Original published cell-type annotations are shown, with endocrine and exocrine populations in grey, and mesenchymal (yellow) and endothelial cells (cyan) highlighted for downstream analysis. (D) Relative enrichment of ECM and BM gene programs across cell types. Dot color represents the relative mean gene set score scaled across cell types for each gene program (standard_scale=’var’). Mesenchymal cells (n = 2,763) exhibit the highest relative enrichment of both ECM and BM programs (relative mean score = 1 for both), followed by ECs (n = 2,097, ECM = 0.310; BM = 0.508). Ductal cells (n = 2,197) displayed minimal ECM enrichment (0.00) with moderate BM enrichment (0.280). Endocrine populations (n = 33,220) exhibited comparatively low enrichment of both programs, with ECM scores (0.235, 0.079, 0.098, 0.159, 0.101) and BM scores (0.092, 0.062, 0.055, 0.073, 0.033) across beta, alpha, delta, PP, and epsilon cells, respectively. (E) UMAP projection of subclustered mesenchymal cells (n = 2,763) reveals two transcriptionally distinct states: MSL1 cells (blue, n = 1,813) and MSL2 cells (red, n = 950). Dot plot of cluster-specific gene expression profiles shows that MSL1 cells are characterized by expression of *S100A6*, *LUM*, and *DCN*, consistent with a fibroblast-like phenotype, whereas MSL2 cells are enriched for expression of mural-associated genes including *RGS5*, *PDGFRB*, and *ADIRF*. (F) UMAP embedding of general ECM and BM gene set scores. MSL2 (pericyte-like) cells exhibit higher BM enrichment relative to ECM (BM mean 0.215 vs ECM mean - 0.007), whereas MSL1 (fibroblast-like) cells display a broader ECM program (ECM mean 0.112 vs BM mean 0.302). Color indicates the relative mean gene set score for each cell type, calculated using sc.tl.score_genes and scaled across cell populations. (G) Divergent matrix expression by mesenchymal subpopulations. Violin plots show the distribution of single-cell log1p-normalized expression of indicated genes. Plot width reflects cell density, central black dashed lines indicate the median, and colored dashed lines represent the interquartile range (IQR). Statistical significance was assessed using a two-sided Mann-Whitney U test. Left, comparison of vascular BM gene expression. Expression of *COL4A1*, a structural BM protein, is significantly higher in MSL1 compared to MSL2 cells (median expression 1.753 vs 1.506, respectively; p < 0.001). However, there is no difference in *COL4A2* expression between the cell types (median expression 1.90 vs 1.797, respectively; not significant (ns)). Right, comparison of interstitial ECM gene expression. Both genes are significantly enriched in fibroblast-like MSL1 cells (*COL1A1* median expression 1.841; *COL1A2* median expression 1.775) and expressed to a significantly lower level in pericyte-like MSL2 cells (median expression 0; p < 0.0001 for both). (H) Heatmap showing mean log1p-normalized expression of published stromal marker gene sets across (top) mesenchymal populations identified in this study (MSL1 and MSL2) and (bottom) previously annotated stromal cell types from public datasets, including Tabula Sapiens (TS)[71] (Pancreas), the Craig-Schapiro dataset[25], and the Human Cell Atlas[72, 73] (Pancreas). Reference cell types include fibroblasts, stellate cells, smooth muscle cells, and pericytes. Heatmap color represents mean gene expression per cell type.

**Figure 2:**
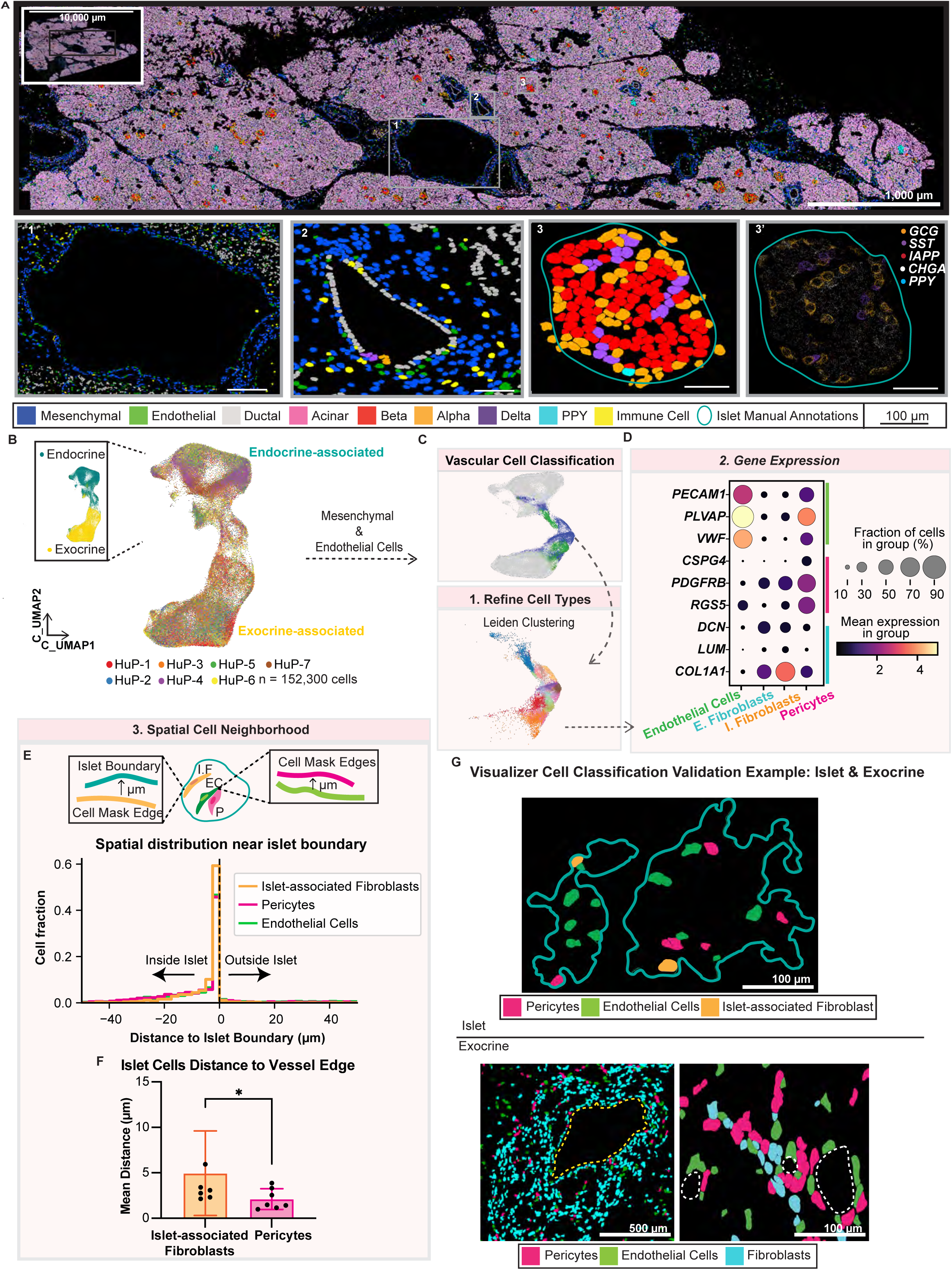
Spatial transcriptomics resolves vascular-associated stromal populations and confirms pericyte identity in the adult human pancreas. (A) Representative MERSCOPE spatial cell-type map of human pancreatic tissue demonstrating preserved endocrine and exocrine architecture. Cell-type annotations projected onto segmentation-derived MERSCOPE cell masks (colors) reveal the spatial organization of endocrine, exocrine, vascular, stromal, and immune populations. Islets were manually delineated based on tissue morphology and enrichment of canonical endocrine transcripts (Inset 4; *GCG*, *CHGA*, *SST*, and *PPY*). Insets highlight the spatial relationships between islets, vascular structures, and pancreatic ducts at increasing magnification. Scale bars = 10,000 µm, 1,000 µm, and 100 µm from top to bottom. (B) UMAP embedding of 152,300 cells, 76,315 exocrine-associated cells (downsampled from 3,270,093 cells) and 75,985 endocrine-associated cells were integrated using CONCORD. Samples span seven human pancreas (HuP) donors. Mesenchymal cells and ECs are subsetted for further analysis. Note that cells cluster first by region and then by type, presumably due to mixing of transcripts between neighboring cells within each region[86, 127]. (C) Vascular cell classification by Leiden clustering and embedding of vascular-associated cells (n = 18,540), resolving 15 transcriptionally distinct subclusters that were subsequently consolidated into endothelial, pericyte, fibroblast, and islet-associated fibroblast populations, distributed across islet-associated (n = 6,217 cells) and exocrine-associated (n = 12,323 cells) anatomical compartments. (D) Annotation of vascular-associated populations. Dot plot shows expression of genes used to define ECs (n = 9,911; *PECAM1*, *VWF*, *PLVAP*), pericytes (n = 3,159; *RGS5*, *PDGFRB*, *CSPG4*), and fibroblast populations (n = 4,739; *DCN*, *LUM*). A subset of islet-associated fibroblast-like cells exhibits low endothelial marker expression and minimal *RGS5*, while retaining fibroblast markers, and was provisionally classified as islet-associated fibroblasts (n = 731). (E) Top, schematic of spatial neighborhood analysis performed by comparing distances between cell boundaries and either the nearest islet boundary or neighboring cell types. Bottom, histogram shows normalized cell fraction as a function of distance to the islet edge, with negative values representing cells within the islet boundary and positive values representing cells outside the islet boundary. Dashed line (x = 0) indicates the islet boundary. (F) Bar plot of mean distance ± SD for each cell group, with each black dot representing an individual donor pancreas. Statistical significance between groups (n = 7 donors) was assessed using Wilcoxon matched-pairs signed rank test. Pericytes (mean distance 2.07 µm) and islet-associated fibroblasts (mean distance 4.97 µm) are located on average 2–5 µm from vasculature, with pericytes significantly closer to ECs (p = 0.016). (G) *In situ* validation of vascular-associated populations by representative MERSCOPE fields of view. Within islets (*top*, teal outline), pericytes (magenta) are tightly associated with ECs (green) along vascular structures, while islet-associated fibroblasts (peach) localize to the islet periphery. In exocrine regions (*bottom*), fibroblast (cyan) populations are more broadly distributed between vascular structures (white dotted outline) composed of ECs and pericytes and enriched around ductal regions (yellow dotted outline). Scale bar = 100 µm.

**Figure 3:**
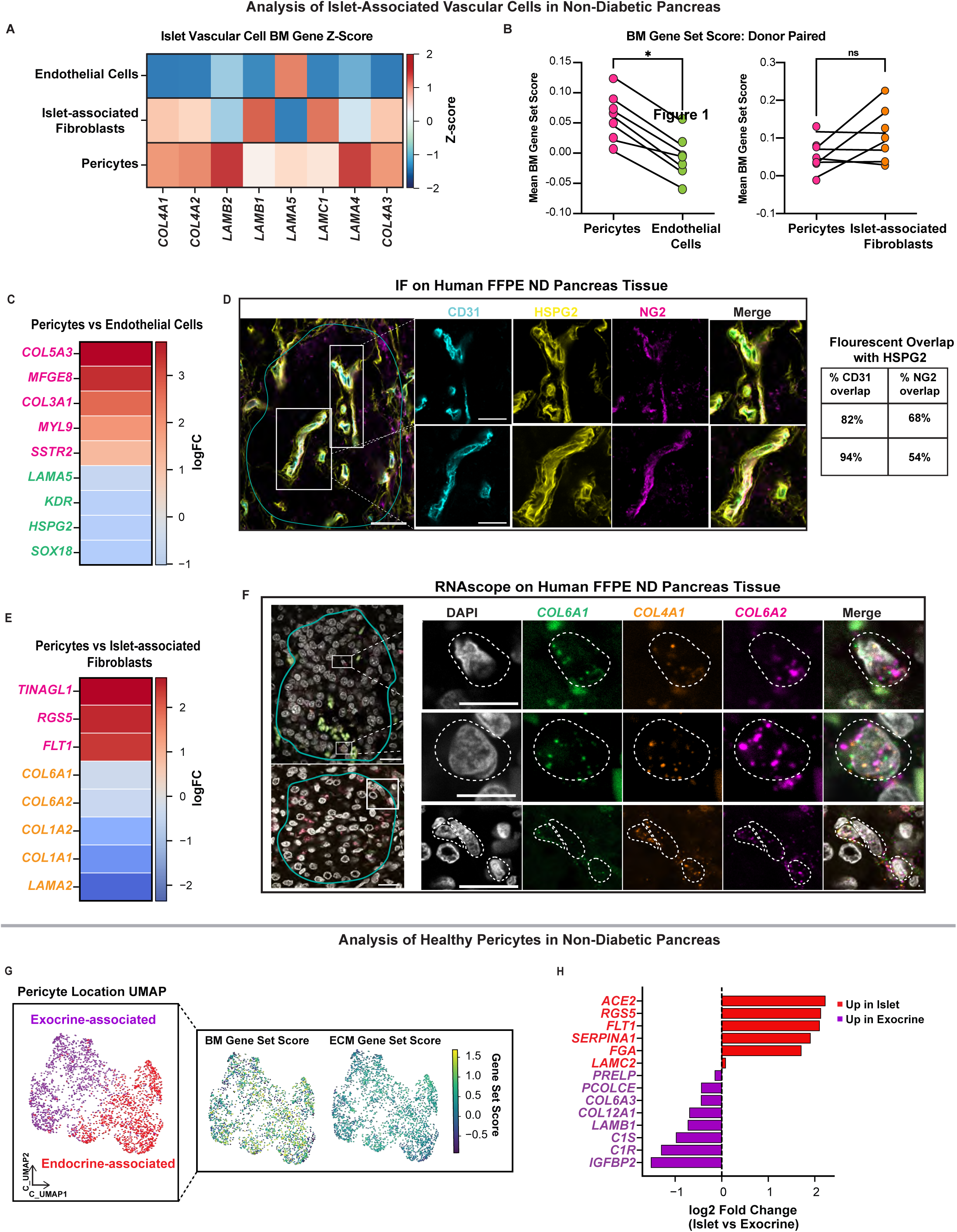
Differential ECM transcriptional programs in the endocrine and exocrine vascular compartment. (A) Heatmap showing z-score normalized expression of curated BM genes across ECs, islet-associated fibroblasts, and pericytes within the islet vascular niche. For each gene, expression values were averaged within each cell type and subsequently z-score scaled across populations to visualize relative enrichment patterns independent of absolute expression magnitude. Positive z-scores indicate relative enrichment within a given population, whereas negative values indicate comparatively lower expression relative to the other populations. (B) Comparison of BM gene-set scores across donor-paired cell populations (n = 7). Statistical significance was assessed via the Mann-Whitney U test. Pericytes have significantly higher BM scores compared to ECs (median 0.015 vs -0.048; p = 0.016). No significant difference exists between pericyte and islet-associated fibroblast scores (median 0.056 vs 0.039; p = 0.468). (C) Differential gene expression (DGE) heatmap comparing pericytes and ECs in the spatial dataset. Color scale represents log2 fold change in expression between populations, with positive values indicating relative enrichment in pericytes (magenta) and negative values indicating relative enrichment in ECs (green). Differentially expressed genes were cross-referenced against independent scRNA-seq reference datasets to minimize potential confounding from spatial segmentation and transcript diffusion artifacts (Fig. S5B). (D) Immunofluorescence image showing the vasculature within a representative ND human islet (outlined in teal), stained for CD31 (endothelial; cyan), HSPG2 (BM; yellow), and NG2 (pericyte; magenta). Right, quantitative channel colocalization analysis performed using ImageJ Coloc2 demonstrating stronger spatial association between endothelial and BM-associated HSPG2 signals within the intra-islet vasculature. 63x; scale bars = 40 µm for large image and 5 µm for higher-magnification images. (E) DGE heatmap comparing pericytes and islet-associated fibroblasts from the spatial dataset. Color scale represents log2 fold change in expression between populations, with positive values indicating relative enrichment in pericytes (magenta) and negative values indicating relative enrichment in islet-associated fibroblasts (orange). Differentially expressed genes were validated against independent scRNA-seq reference datasets to reduce potential bias introduced by segmentation or transcript diffusion artifacts (Fig. S5C). (F) Multiplexed RNA *in situ* hybridization image of ND human pancreatic tissue, showing expression of *COL6A1* (green), *COL4A1* (orange), and *COL6A2* (magenta) transcripts; nuclei are stained with DAPI in white. Representative positive cells (outlined in dashed white lines) were selected for inspection at higher magnification to verify presence of ECM-expressing cells in various endocrine niches using an orthogonal method to detect RNA. 63x; scale bars: 40 µm (top), 20 µm (bottom) in large image and 5 µm in higher-magnification insets. (G) (*left*) UMAP embedding of pericytes from spatial dataset reveals segregated clusters based on anatomical location: endocrine-associated (red) versus exocrine-associated (purple) populations. (*right*) Spatial projection of matrix programs onto the pericyte UMAP embedding. Color represents individual cell gene set score. (H) Differential expression analysis highlighting transcriptional differences between islet-and exocrine-associated pericytes. Bars represent log2 fold change in expression between islet and exocrine compartments, with positive values indicating relative enrichment in endocrine-associated pericytes (red) and negative values indicating relative enrichment in exocrine-associated pericytes (purple). Differentially expressed genes were cross-referenced against independent scRNA-seq reference datasets to minimize potential confounding from spatial segmentation and transcript diffusion artifacts (Fig. S5D).

### Differential Expression Analysis

Differential gene expression (DGE) analysis was performed using Scanpy’s sc.tl.rank_genes_groups with the Wilcoxon rank-sum test. Analyses were conducted on log-transformed expression values (use_raw = False), and p-values were adjusted using the Benjamini-Hochberg procedure to control the false discovery rate.

### RNA DV200 Analysis

To assess RNA integrity prior to MERFISH analysis, RNA was extracted from FFPE tissue using the Zymo Quick-RNA FFPE Miniprep kit (Zymo Research, R1008; RRID:SCR_024103). RNA concentration was measured by NanoDrop (RRID:SCR_018042) spectrophotometry. RNA fragment size distribution and DV200 values were assessed using the Agilent 4200 TapeStation (RRID:SCR_018435) RNA ScreenTape assay according to the manufacturer’s protocol. DV200 values represent the percentage of RNA fragments longer than 200 nucleotides.

### Gene selection for MERFISH

A custom 300-gene MERFISH panel was designed using the Vizgen MERSCOPE Gene Panel Design Portal (https://portal.vizgen.com) [58]. Gene selection was informed by integration of published human single-cell/nucleus RNA-seq datasets, proteomic studies of human pancreas, Human Protein Atlas (RRID:SCR_006710), and the CellxGene tool. Genes were selected based on probe design feasibility (sufficient probe binding sites), expression abundance thresholds (FPKM), and avoidance of optical crowding during imaging. Insulin (*INS*) was excluded due to probe design constraints related to transcript abundance and sequence composition. The final 300-gene panel included 3 categories. Category 1 genes comprised 98 ECM genes reported to be expressed in the human pancreas and differentially expressed in islets and exocrine tissue at the protein level. Category 2 genes were composed of 25 canonical human endocrine, exocrine, and vascular cell-type markers widely used in the field to annotate known cell types (e.g., *GCG* to mark alpha cells, *PECAM1* to mark endothelial cells, *KRT19* to mark ductal cells). Category 3 genes were reported in recently published literature as being expressed by subpopulations of beta cells and endothelial cells in the pancreas. To serve as a control for nonspecific binding of probes, we included 15 blank barcodes. The full gene panel csv file is available in Table S3.

### MERFISH for FFPE tissue samples

A fully detailed, step-by-step guide to the MERFISH sample preparation full protocol is available at https://vizgen.com/resources/fresh-and-fixed-frozen-tissue-sample-preparation. Briefly, FFPE samples from selected donors were sectioned using a microtome (Leica Microtome, HM325; RRID:SCR_020259) at 4-µm thickness and placed on a MERSCOPE FFPE slide (Vizgen, catalog no. 20400100) in accordance with Vizgen’s MERSCOPE User Guide for FFPE samples. Tissues were cleared using the resistant-clearing protocol for the allowed maximum time (72 hours). After tissue clearing, samples were treated with MERSCOPE Photobleacher (Vizgen, catalog no. 10100003) for 4 h, washed with Formamide Wash Buffer at 37 °C for 30 min, and then incubated with MERSCOPE Gene Panel Mix for 48 hours at 37 °C.

### Cell Segmentation

Automated cell segmentation was performed on nuclear and cell membrane stains using Cellpose v2.0 (RRID:SCR_021716) within the MERSCOPE analysis workflow[59]. Segmentation outputs included cell masks and associated cellular metadata stored in .json files. In addition, stitched mosaic images corresponding to the DAPI, cell boundary, and poly(A) channels were exported as .jpeg files.

### Islet Annotations

Manual islet annotation was performed in MERSCOPE Visualizer (2.1.2593.1). Annotations were selected using four criteria: (1) expression of endocrine-associated transcripts within the candidate region, (2) presence of DAPI signal confirming cellular localization, (3) presence of cells annotated as endocrine populations based on transferred cell labels displayed in the Visualizer overlay, and (4) a minimum of three endocrine cells per annotated islet. Regions meeting all criteria were annotated as islets. Annotated islet geometries and associated metadata, including transcript coordinates, cell identities, and islet assignments for each cell were exported as .csv files for downstream analyses (Tables S8-S9). A total of 2,387 ND islets and 1,311 T2D islets were annotated.

### Spatial Transcriptomics Quality Control and Filtering

Cells with fewer than 10 detected transcripts were removed during quality control. No genes were excluded from the dataset, and all 300 genes included in the MERFISH panel were retained for downstream analyses. Raw counts were retained in adata.raw and adata.layers[‘counts’]. For downstream analyses, transcript counts were normalized by total counts per cell using Scanpy’s normalize_total function and subsequently log-transformed using log(1 + x).

### Spatial Transcriptomics Clustering and Cell-type Annotation

Spatial cell-by-gene expression matrices generated from the Vizgen MERSCOPE platform were analyzed in Python using Scanpy and Squidpy. Raw transcript counts were normalized and log-transformed (log(1 + x)). To resolve vascular and mesenchymal heterogeneity, cells initially annotated as mesenchymal or endothelial were isolated from the full dataset and reclustered using Leiden clustering (resolution = 1.0). Differential expression analysis was repeated on the resulting clusters, and the top 20 marker genes were used to guide cluster annotation.

### Spatial Distance Calculations

Spatial relationships between vascular-associated populations were quantified using segmentation-derived cell polygons and Shapely geometric operations. Cell boundaries generated from MERSCOPE segmentation masks were converted into polygon geometries and indexed using STRtree spatial indexing. Because polygon coordinates are reported in the native MERSCOPE coordinate system (microns), all distances are reported in microns. For cell-to-cell analyses, the shortest edge-to-edge distance between cell boundaries was calculated using Shapely’s .distance() method applied to polygon geometries. Distances between ECs, pericytes, fibroblasts, islet-associated fibroblasts, and endocrine cells were determined using nearest-neighbor searches across annotated cell populations. For islet boundary analyses, manually annotated islet polygons generated in MERSCOPE Visualizer were used to define endocrine compartment boundaries. The shortest edge-to-edge distance between each cell polygon and the nearest islet boundary polygon was similarly calculated using Shapely’s .distance() method. Cells located within the annotated islet region were assigned negative distance values, whereas cells located outside the islet boundary were assigned positive values. Distances were summarized at both the cell level and donor level for downstream statistical analyses.

### Cell Composition Analysis

Vascular-associated cell composition was assessed using annotated endothelial cells, pericytes, fibroblasts, and islet-associated fibroblasts identified from the integrated spatial transcriptomics dataset. For each donor, the total number of cells belonging to each vascular-associated population was quantified and expressed as a proportion of all vascular-associated cells within the analyzed region (islet or exocrine). Cell composition was calculated on a donor-by-donor basis prior to statistical testing. Endothelial-to-pericyte ratios were calculated by dividing the number of islet endothelial cells by the number of islet pericytes identified within each donor. Islet vascularization was calculated as the ratio of endothelial cells plus pericytes to the total number of endocrine cells within each donor. Similarly, the relative abundance of pericytes was calculated as the percentage of pericytes divided by total islet vascular-associated cells. For disease-state comparisons, donor-level composition metrics were compared between non-diabetic and type 2 diabetic samples using Mann-Whitney U tests.

### Immunofluorescence (IF) Staining of Human FFPE Pancreatic Tissue

FFPE human pancreatic tissue sections (4–5 µm thick) were baked at 55°C for 15 minutes prior to processing. Sections were deparaffinized in xylene and rehydrated through graded ethanol washes to distilled water. Antigen retrieval was performed using heat-mediated retrieval in citrate buffer pH = 6.0 (Vector Labs, H-3300-250) for 15 minutes and then cooled to RT for another 15 minutes. Tissue sections were permeabilized and blocked in 0.5% PBST with 5% normal donkey serum (Jackson ImmunoResearch Labs Cat# 017-000-121, RRID:AB_2337258) for 1 hour at RT. Primary antibodies were incubated overnight at 4 °C in a humidified chamber. Following three 1X PBS washes, sections were incubated with species-specific AlexaFluor-conjugated secondary antibodies at RT for 1 hour. Nuclei were counterstained with DAPI if needed. Slides were mounted using antifade mounting media (Vector Labs, H-1000-10, RRID:AB_2336789). Fluorescence imaging was performed using a Zeiss LSM800 laser-scanning confocal microscope (RRID:SCR_015963). For colocalization analysis, images were processed in Fiji/ImageJ and quantified using the Coloc2 plugin[60]. Mean fluorescence intensity was quantified within manually annotated CHGA-defined islet regions in Fiji using identical processing, staining, incubation, and image acquisition settings for all samples. A complete list of antibodies, vendors, catalog numbers, working dilutions, and RRID’s is provided in Supplementary Table S22.

### RNAscope Multiplex Fluorescence Assay on Human FFPE Pancreatic Tissue

RNAscope multiplex fluorescence *in situ* hybridization was performed on FFPE human pancreatic tissue sections using the RNAscope Multiplex Fluorescent Reagent Kit (Advanced Cell Diagnostics, Cat. No. 323100 & 323270) according to the manufacturer’s protocol[61]. Target-specific RNAscope probes were hybridized to tissue sections followed by sequential signal amplification steps according to the manufacturer’s instructions. Fluorescent detection was performed using Opal fluorophores (Akoya Biosciences). Nuclei were counterstained with DAPI, and slides were mounted using an antifade mounting medium (Vector Labs, H-1000-10). Fluorescence imaging was performed using a Zeiss LSM800 laser-scanning confocal microscope at either 40x or 63x magnification. Table of probes used is provided in SI (Table S22).

### Gene set enrichment analysis

We performed gene set enrichment analysis (GSEA) separately for MSL1 (fibroblast-like) and MSL2 (pericyte-like) populations using GSEApy.

### Identification of Islet-Associated Fibroblasts

A distinct islet-associated fibroblast population was identified in the spatial transcriptomics data that was observed to localize preferentially to the islet boundary and exhibited elevated expression of fibrillar collagen genes (*COL1A1*, *COL1A2*, and *COL3A1*), basement membrane-associated genes (*LAMA2*, *LAMB1*, *LAMC1*, and *LAMC3*), and ECM-remodeling genes including *PRELP*, *FBLN1*, and *COL6A3*. These cells were annotated as islet-associated fibroblasts (IAFs). Independent validation was subsequently performed using the PancDB and Craig-Schapiro human pancreas reference datasets.

### Cross-Reference Validation with External scRNA-seq Datasets

Comparisons were performed against external human pancreas scRNA-seq reference datasets PancDB and the Craig-Schapiro pancreas dataset. Spatial populations were compared against reference populations using shared genes present across datasets and restricted to genes included within the spatial transcriptomics panel where indicated. For transcriptional concordance analyses, average gene expression profiles were calculated for each annotated spatial and reference population. Pearson correlation and Spearman correlation metrics were used to quantify cross-platform similarity between matched populations. Gene-level comparisons were performed using normalized mean expression values across shared genes. Differentially expressed genes identified in spatial analyses were additionally cross-referenced against reference datasets. Where PancDB lacked explicit anatomical annotations, corresponding stromal populations were identified computationally based on transcriptional similarity and marker gene enrichment.

### Statistical Analysis

Statistical analyses were performed in Python using SciPy and in GraphPad Prism (v10.6.1; RRID:SCR_002798). For donor-level analyses, paired comparisons were assessed using two-sided Wilcoxon matched-pairs signed-rank tests and unpaired comparisons between non-diabetic and type 2 diabetic donors were assessed using two-sided Mann-Whitney U tests. Cell-level comparisons were performed using two-sided Mann-Whitney U tests. Donors were considered the biological replicate for donor-level analyses, whereas individual cells were used as the unit of analysis for cell-level comparisons. Non-parametric statistical tests were selected because they do not require assumptions of normality and are appropriate for the sample sizes used in this study. Data are presented as indicated in the figure legends. P values < 0.05 were considered statistically significant.

**Additional experimental and computational methods are described in the Electronic Supplementary Material (ESM), Supplementary Materials and Methods.**

## Results

### Mesenchymal cells are the primary source of vascular basement membrane gene expression in the healthy adult human pancreas

Previous studies in the mouse have indicated a central role for the vascular compartment in forming the islet BM[17, 62, 63]. As the vascular BM in human islets differs in its uniquely bilayered structure (Fig. 1A), we set out to better understand the cellular sources and composition of human islet vascular BM. To gain an unbiased understanding of the gross composition and BM gene expression patterns across cell types in the human pancreas, we first re-integrated publicly available scRNA-seq data from PancDB, restricting our analysis to non-diabetic adult donors (n = 25 donors; n = 88,701 cells)[57] (Fig. 1B). Three samples were removed because of low quality RNA (see Methods; Table S1). We performed additional quality control, doublet filtering, and batch alignment (Fig. S1A-C) with CONCORD, a recently reported tool for data integration, dimensionality reduction, and denoising that preserves biological features better than other reported methods[55] (Fig. S1D).

Following integration and clustering, cells from multiple batches co-embedded into well-resolved clusters representing all major pancreatic cell populations (n = 22 donors, n = 46,106 cells; Fig. 1C). To assess BM and ECM gene expression across pancreatic cell types, we calculated gene set scores (Fig. 1D) using the NABA Basement Membrane[64, 65] and NABA Core Matrisome[65, 66] gene sets obtained from Molecular Signature Database (MSigDB)[67, 68]. To distinguish broader ECM programs from BM-associated genes, genes present in the BM set were excluded from the Core Matrisome set prior to scoring. The resulting gene lists are provided in Table S2. Surprisingly, this analysis revealed that mesenchymal cells (relative mean score = 1), rather than ECs (0.508), exhibited the highest relative enrichment of BM scores compared to other cells. In contrast, endocrine cells exhibited substantially lower ECM and BM gene program enrichment, consistent with previous reports in the mouse[13]. Although ductal cells showed a positive BM signal (0.280), this enrichment was restricted to a subset of cells expressing modest levels of BM-associated genes, most notably *COL18A1* and laminin subunits (Fig. S1E, Table S3).

Given the enrichment of BM gene expression within the mesenchymal population, we next examined heterogeneity within this compartment. Unsupervised sub-clustering revealed two transcriptionally distinct populations (Fig. 1E, left; Table S4). The first population, mesenchymal subtype-1 (MSL1), expressed fibroblast-associated genes, including *S100A6* and *LUM*, indicative of a fibroblast-like identity[69, 70]. The second population, mesenchymal subtype-2 (MSL2), was enriched for mural cell genes, such as *RGS5*, *PDGFRB*, and *C11orf96*, consistent with a pericyte-like identity (Fig. 1E, right; S1F-J)[25]. Although both populations expressed ECM genes, MSL2 showed preferential enrichment of BM programs, with a higher mean BM gene set score (0.215) than ECM score (−0.007). MSL1 cells also exhibited elevated BM-associated gene expression but showed concurrent enrichment of broader ECM programs, with BM and ECM scores of 0.302 and 0.112, respectively (Fig. 1F).

To further resolve the differences in BM gene expression between MSL1 and MSL2 populations, we compared the expression of core BM genes (*COL4A1*, *COL4A2*) and interstitial collagens (*COL1A1*, *COL1A2*) (Fig. 1G). Fibroblast-like MSL1 cells displayed significantly higher expression of *COL4A1* (p < 0.001) (Fig. 1G, left), while the two populations expressed indistinguishable levels of *COL4A2*. Interstitial collagens *COL1A1* and *COL1A2* were significantly enriched in fibroblast-like MSL1 cells and largely absent in MSL2 (p < 0.001) (Fig. 1G, right). These results indicate that MSL1 fibroblast-like cells express a mixed ECM program, while MSL2 pericyte-like cells are more specialized for BM gene expression.

To more precisely classify the MSL1 and MSL2 populations, we compared their transcriptional profiles against established mesenchymal-type signatures from Tabula Sapiens[71], the Craig-Schapiro dataset[25], and the Human Cell Atlas[72, 73] (Fig. 1H, S1K). This analysis revealed two recurring obstacles in defining pancreatic mesenchymal identity through transcriptomics alone. First, the marker genes historically used to distinguish fibroblasts, stellate cells, smooth muscle cells (SMCs), and pericytes overlap substantially (Fig. S1L). Genes such as *PDGFRB*, *ADIRF*, and *CSPG4*, for example, are broadly expressed across multiple clusters rather than being restricted to pericytes as frequently assumed. Second, this marker promiscuity is reflected in the inconsistent labeling of transcriptionally similar populations across datasets. For instance, cells annotated as “pericytes” in one study share expression profiles with cells called “stellate cells” in another, and stellate cells and fibroblasts are often indistinguishable by gene expression alone. These inconsistencies reflect that pancreatic mesenchymal populations may share developmental origins and exhibit functional plasticity, producing transcriptional overlap that confounds annotation [69, 74–76]. Taken together, resolving pancreatic mesenchymal populations into defined cell types like pericytes or fibroblasts requires additional spatial information alongside transcriptional profiling [75, 77].

### Spatial transcriptomics identifies and maps pericyte, endothelial, and fibroblast populations within the human pancreas

To definitively resolve the identity of the BM-producing cells in the islet as either pericytes or fibroblasts, we turned to spatial transcriptomics. Specifically, we aimed to define pericytes based on: (1) expression of a mural gene program (*PDGFRB*, *CSPG4*, *RGS5*) and (2) physical proximity to ECs. This strategy additionally enabled us to build a framework to understand how BM gene programs are spatially organized within the endothelial and perivascular compartments of the pancreatic vasculature. We achieved this by performing MERFISH spatial genomics on adult human pancreas tissue sections from seven non-diabetic individuals using the Vizgen MERSCOPE platform. Donor demographic and clinical information are summarized in Table 1 and detailed in Table S5. We prepared a targeted 300-gene panel designed to annotate all major pancreatic cell types and interrogate the expression of ECM genes. Gene selection was informed by manual curation of publicly available human pancreatic RNA-sequencing datasets, human pancreatic proteomic studies, the CellxGene platform, and the Human Protein Atlas (Fig. S2A, see Methods)[70, 71, 78–81]. The final probe panel included canonical markers for endocrine, exocrine, immune, endothelial, and mesenchymal populations, as well as 98 ECM-associated genes enriched for BM and interstitial ECM components that are expressed within the pancreas[4, 80, 82] (Table S6).

Whole-section spatial maps revealed that both islets and surrounding exocrine tissue were intact and displayed the expected histological architecture (Fig. 2A, top; representative sample shown). Mapping annotated cell identities back to their spatial coordinates revealed the full diversity of the pancreatic cellular landscape, including endocrine, exocrine, vascular, and immune populations (Fig. S2B) [83]. To enable compartment-specific analysis, we manually segmented 2,387 islet boundaries based on cell morphology and the localized enrichment of canonical endocrine transcripts (*GCG*, *CHGA*, *SST*, *PPY*, *IAPP*) (Table S7). Higher magnification of selected regions further validated the precise spatial organization of vascular and ductal networks, providing a high-confidence dataset for investigating the vascular niches present in the adult pancreas (Fig. 2A, bottom).

To generate a unified representation across donors, we next integrated all spatial datasets using CONCORD (Fig. S2C). From 3,270,093 segmented cells, we retained all islet cells and down-sampled exocrine cells to a similar magnitude, producing a final embedding of 152,300 cells across two major clusters: endocrine-annotated (teal, n = 75,985) and exocrine-annotated (yellow, n = 76,315) (Fig. 2B, S2D). Major pancreatic cell types formed resolved clusters, with expected gene expression patterns, including endocrine cells expressing *CHGA* and *GCG*, immune cells expressing *CD8A*, and ductal cells expressing *KRT19* [73, 78, 83, 84](Fig. S2E-F).

To further refine vascular-associated populations, we restricted downstream analysis to the mesenchymal and endothelial populations (Fig. 2C). Unsupervised clustering identified multiple transcriptionally distinct mesenchymal and vascular populations, which were annotated using differential gene expression (DGE) analysis together with analysis of canonical lineage marker gene expression (Fig. 2D, S2G). ECs were identified by expression of *PECAM1*, *VWF*, and *PLVAP*, whereas pericytes were defined by enrichment of *RGS5*, *PDGFRB*, and *CSPG4* together with additional vascular-associated transcriptional features[25, 73]. Fibroblast populations were characterized by expression of *DCN* and *LUM* (Fig. S2H). Unsupervised clustering revealed two transcriptionally distinct fibroblast populations. One population comprised most fibroblasts and was localized predominantly throughout the exocrine pancreas. In contrast, a second fibroblast cluster retained canonical fibroblast markers while exhibiting enrichment of ECM genes, including *COL1A1*, *LAMA2*, *COL1A2*, *COL5A1*, and *COL3A1* (Fig. S2I). This population also showed detectable endocrine transcripts (*GCG*, *CHGA*), likely reflecting transcript diffusion or contamination from neighboring endocrine cells due to its close spatial association with islets. Together, these features suggested the presence of a specialized fibroblast population associated with the islet microenvironment, which we designated islet-associated fibroblasts, whereas the broader fibroblast population was designated as exocrine-associated fibroblasts (Fig. S2J). To validate this population, we initially compared fibroblast populations against the PancDB human pancreas scRNA-seq atlas, which lacks predefined spatial annotations and is free from contamination artifacts unique to spatial transcriptomics data such as transcript diffusion and segmentation errors (Fig. S3)[70, 85, 86]. Sub-clustering of the MSL1 (fibroblast-like) population described in Figure 1 identified a fibroblast subset characterized by enrichment of *COL1A1*, *COL1A2*, *COL3A1*, *SPARC*, *COL4A1*, *COL5A2*, *COL5A1*, and *LGALS1* (Fig. S3B-F). Notably, this ECM-rich signature distinguished the population from other fibroblast subsets in PancDB and closely matched the transcriptional profile of the spatially identified islet-associated fibroblasts (Fig. S3A). To further validate the identity of the islet-associated fibroblast population, we compared average gene expression profiles to the location-resolved Craig-Schapiro dataset using Pearson and Spearman correlation analyses (Fig. S3G)[87]. Spatial islet-associated fibroblasts showed high similarity to annotated islet fibroblasts across both metrics (Pearson r = 0.608; Spearman ρ = 0.664), providing independent support for the anatomical assignment and transcriptional distinctiveness of this endocrine niche-associated fibroblast population (Fig. S3H).

To validate these classifications spatially, we next quantified spatial neighborhood relationships at both the cell and donor level. Consistent with preliminary annotations, cells labeled as islet-associated fibroblasts (67%) localized preferentially within ± 5 µm of the islet boundary across donors (Fig. 2E, S2K-L). We further validated annotations by measuring the distance from each cell boundary to the vasculature, using the nearest EC mask edge as a proxy for the vessel lumen. Both pericytes and islet-associated fibroblasts were consistently located within 5 µm of vasculature, with pericytes significantly closer than islet-associated fibroblasts (p = 0.016) to ECs (Fig. 2F, S2M-N). We confirmed that within islets, the cells annotated as pericytes were tightly associated with ECs along vascular structures, whereas islet-associated fibroblasts were enriched at the islet periphery (Fig. 2G; top). In exocrine regions, exocrine-associated fibroblasts were predominantly organized within the interstitial stroma, sequestered between periductal bands (yellow dotted lines) and vessel-surrounding stroma (Fig. 2G; bottom, S2O). Notably, exocrine-associated fibroblasts remained spatially excluded from the immediate perivascular niche, which was predominantly occupied by pericytes in direct contact with the endothelium (white dotted outlines). Together, these analyses confirm that the mesenchymal population annotated as pericytes by scRNA-seq corresponds to bona fide vascular-associated cells *in situ* and occupies the immediate perivascular niche within islets. Importantly, accurate annotation of these populations was only possible through the integration of manual islet annotation and spatial single-cell transcriptomics.

### Distinct features of ECM gene expression programs in the endocrine versus exocrine vascular compartments

Having validated the identity of endothelial and mural populations within the human pancreas, we next turned to investigating how BM and other ECM gene expression programs are organized within the pancreatic vascular niche. Spatial mapping of *COL4A1*, *COL4A2*, and *LAMC1* transcripts revealed localization along islet vascular structures, with signal detected proximal to vascular-associated cells rather than uniformly distributed across the tissue (Fig. S4A). To quantify these patterns, we examined the expression of a curated set of islet vascular BM genes (*COL4A1*, *COL4A2*, *COL4A3*, *LAMB2*, *LAMB1*, *LAMA5*, *LAMC1*, *LAMA4*) that have been reported to be present at the protein level either in the intra-islet vascular BM, peri-islet BM, or both[4](Fig. 3A). Pericytes displayed the strongest relative enrichment of these core BM components, particularly collagen IV (*COL4A1*, *COL4A2*), together with the laminin subunits *LAMA4* and *LAMB2*. In contrast, ECs exhibited comparatively weaker BM-associated transcriptional signatures within the islet vascular niche, with selective enrichment of *LAMA5*. Islet-associated fibroblasts demonstrated enrichment of a distinct set of laminin subunits, including *LAMB1* and *LAMC1*.

To determine whether these BM transcriptional differences were consistently maintained across donors rather than driven by cell abundance or sampling effects, we performed donor-paired comparisons. BM gene set scores were calculated separately within each donor, allowing direct comparison of matched cell types within each tissue section. Across all seven donors, ECs consistently showed significantly lower BM scores than pericytes (n = 7 donors; median score = 0.015 vs -0.048; p = 0.016; Fig. 3B, left). In contrast, islet-associated fibroblasts had comparable BM scores to pericytes (p = 0.468; Fig. 3B, right).

Differential expression (DE) analysis was next performed to further characterize the broader transcriptional differences between pericytes, ECs, and islet-associated fibroblasts (Fig. S4B-C, Table S9-10). Relative to pericytes, ECs exhibited increased expression of *HSPG2* and *LAMA5*, indicating that ECs contribute directly to vascular BM assembly (Fig. 3C, left)[88, 89]. IF staining confirmed strong spatial association of HSPG2 with CD31+ endothelial structures relative to adjacent pericytes (NG2+) within islets (Fig. 3D). In contrast, islet-associated fibroblasts showed enrichment of a broader ECM program defined by interstitial and fibrillar collagens including *COL6A1*, *COL1A1*, and *COL1A2* (Fig. 3E), reflecting a matrix-producing phenotype distinct from the vascular BM[19]. Multiplexed RNA *in situ* hybridization analysis further identified *COL4A1*+, *COL6A1*+, *COL6A2*+ cells along the islet boundary and occasionally within the intra-islet interior (Fig. 3F); however, we could not exclude the possibility that these interior cells were artifacts of 2D segmentation or sectioning.

Lastly, to investigate whether pericyte transcriptional programs were conserved between the endocrine and exocrine compartments, we subsetted pericytes by location (Fig. 3G, left). Qualitative projection of gene set scores onto the subsetted UMAP suggested that BM and ECM transcriptional activity was broadly consistent across both compartments, with gene set scores distributed relatively uniformly across the UMAP (Fig. 3G, right). To better resolve potential differences, we performed donor-paired analysis. This revealed no significant difference in BM or ECM gene set scores in islet-associated pericytes relative to their exocrine counterparts (Fig. S4D). DE analysis did, however, identify significant niche-specific tuning of broader ECM-related gene programs (Fig. 3H, S4E, Table S11). Exocrine-associated pericytes were enriched for genes linked to interstitial matrix remodeling and inflammatory signaling, including *IGFBP2*, *C1R*, *C1S*, *COL12A1*, *COL6A1*[11, 90–93]. In contrast, islet pericytes exhibited enrichment of *ACE2*, *RGS5*, *FLT1*, *SERPINA1*, *FGA*, and *LAMC2*[94, 95]. These findings indicate that while pancreatic pericytes maintain a conserved core cell identity and vascular maintenance program, they also undergo substantial niche-dependent specialization. Importantly, these niche-associated transcriptional programs were independently supported through analysis of the Craig-Schapiro dataset, where similar compartment-specific expression patterns were observed within the islet and exocrine regions (Fig. S5A-F, Table S12-13).

### Altered Vascular Composition and Spatial Remodeling in Type 2 Diabetic Islets

To investigate how vascular stromal programs are remodeled in type 2 diabetes, we again leveraged the CONCORD re-integrated PancDB atlas to map MSL1 (fibroblast) and MSL2 (pericyte) populations across non-diabetic and type 2 diabetic donors (Figs. 4A, S6A-C; Table S14-15). Although both populations remained identifiable across disease states, gene set enrichment analysis (GSEA) revealed opposing regulation of ECM-associated pathways between the two populations in type 2 diabetes (Fig. S6D). Fibroblast-associated populations exhibited reduced enrichment of collagen formation, matrix crosslinking, and fibril assembly pathways, whereas pericyte-associated populations showed increased enrichment of these same pathways, suggestive of a less differentiated pericyte state and a shift toward a more fibrotic phenotype.

**Figure 4:**
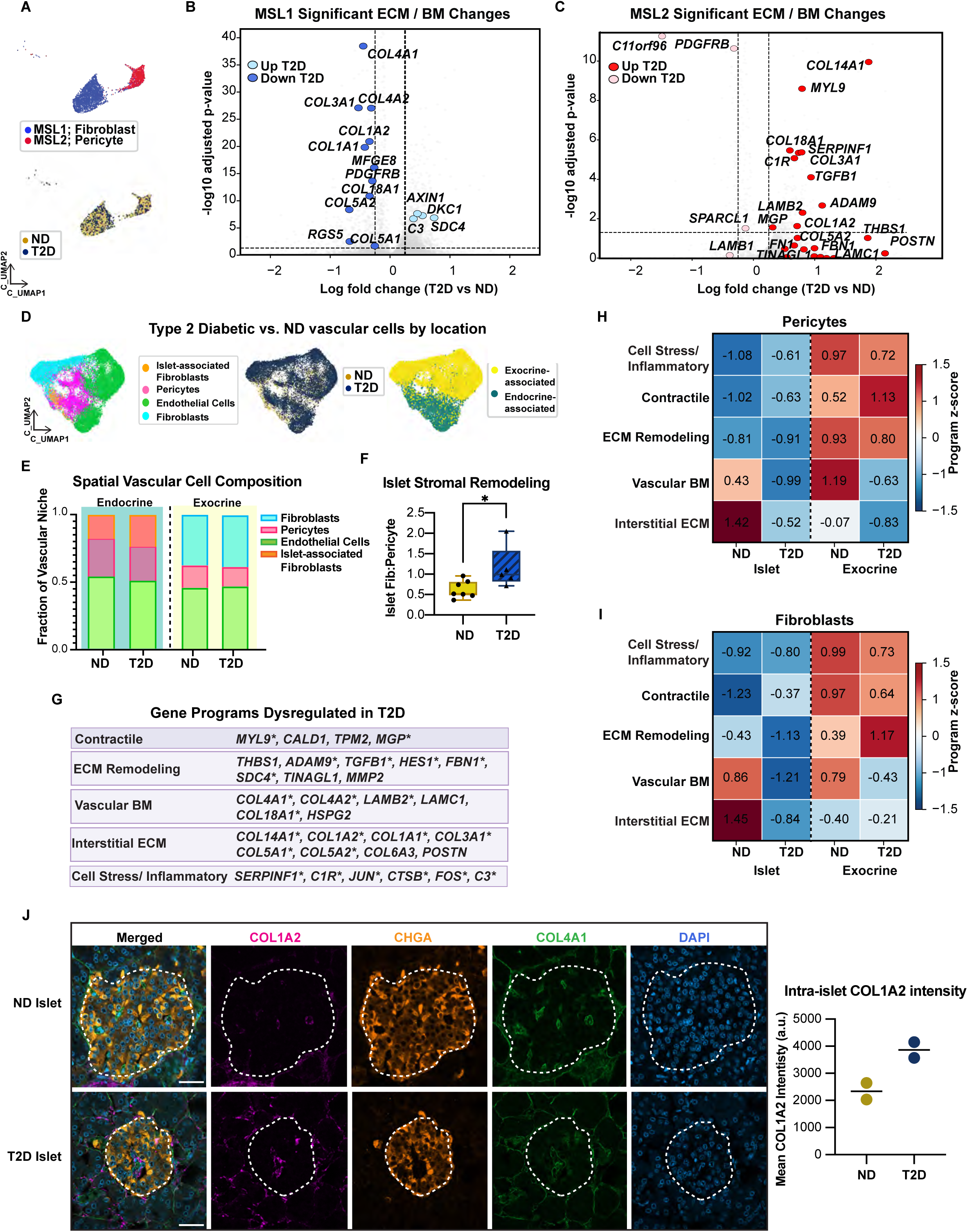
Dysregulation of BM and ECM programs in the Type 2 diabetic perivascular niche. (A) UMAP embedding of PancDB stromal populations including MSL1 fibroblast-like cells (blue) and MSL2 pericyte-like cells (red) across ND and type 2 diabetic (T2D) donors. Bottom panel shows ND (gold) and T2D (navy) sample identities projected into the same embedding. (B) Volcano plot showing ND versus T2D differential expression in MSL1 cells, restricted to genes included in the 300-gene MERSCOPE spatial panel. Light blue points indicate genes significantly upregulated in T2D MSL1 cells, and dark blue points represent genes significantly downregulated in T2D MSL1 cells. Dashed lines indicate log2 fold change and adjusted p-value thresholds used for significance. (C) Volcano plot showing ND versus T2D differential expression in MSL2 cells, restricted to genes included in the 300-gene MERSCOPE spatial panel. Dark red points indicate genes significantly upregulated in T2D MSL2 cells, and light red points represent genes significantly downregulated in T2D MSL2 cells. Dashed lines indicate log2 fold change and adjusted p-value thresholds used for significance. (D) Integrated UMAP embedding of spatial data subsetted for vascular-associated populations from both ND and T2D donors, colored by cell type (left), disease status (middle), and anatomical location (right). Cell populations include ECs, pericytes, fibroblasts, and islet-associated fibroblasts distributed across endocrine (teal) and exocrine (yellow) compartments. (E) Compositional analysis of vascular-associated populations within islet and exocrine compartments in ND and T2D samples. Stacked bars represent the relative fraction of ECs, pericytes, fibroblasts, and islet-associated fibroblasts within the combined vascular-associated compartment for each anatomical region and disease state. (F) Bar plot showing the islet-associated fibroblast-to-pericyte ratio in ND and T2D islets, with each black dot representing an individual donor. Statistical significance between groups (n = 12 donors total) was assessed using an unpaired Mann-Whitney U test. T2D islets exhibited a significantly higher islet-associated fibroblast-to-pericyte ratio (median = 1.38) compared to ND islets (median = 0.90; p = 0.030). (G) Gene programs derived from differential expression analysis between ND and T2D PancDB vascular-associated populations, including Contractile, ECM Remodeling, Vascular BM, Interstitial ECM, Cell Stress/Inflammatory programs. Representative genes contributing to each transcriptional program are shown. (*) Asterisks represent genes significantly differentially expressed in T2D. (H-I) Heatmap showing z-score normalized enrichment of transcriptional programs across spatial pericyte populations (H) or spatial fibroblast populations (I) stratified by disease state and anatomical location. Program scores were calculated independently for each condition and subsequently scaled by z-score across groups to visualize relative enrichment patterns. (J) (*Left*) Representative immunofluorescence images of COL1A2, CHGA, COL4A1, and DAPI staining in ND and T2D human pancreatic islets. Dashed lines indicate CHGA-defined islet boundaries used for quantification. Images were acquired at 63x magnification. Scale bar = 40 µm. (*Right*) Mean COL1A2 fluorescence intensity (arbitrary units, a.u.) was quantified within CHGA-defined islet regions. Each point represents the mean COL1A2 fluorescence intensity of an individual donor; 30 islets were analyzed from 2 ND and 2 T2D donors. COL1A2 fluorescence intensity appeared higher in islets from T2D donors than in ND controls.

To further characterize these transcriptional changes, we performed genome-wide DGE analysis between non-diabetic and type 2 diabetic populations (Fig. S6E-F). Differentially expressed genes included in the spatial transcriptomics panel are shown in the volcano plots in Fig. 4B-C. Fibroblast-associated populations exhibited reduced expression of several BM-associated collagens, including *COL4A1* and *COL4A2* (Fig. 4B). In contrast, pericyte populations exhibited increased expression of contractile, fibrillar ECM, and angiogenic-related genes including *MYL9*, *COL1A2*, *COL14A1*, and *SERPINF1*, together with reduced expression of *PDGFRB*, a receptor critical for pericyte survival and perivascular retention (Fig. 4C)[96–99].

We next sought to determine how these transcriptional states were compositionally and spatially organized within intact tissue. Integration of non-diabetic and type 2 diabetic spatial datasets demonstrated preservation of the major vascular and stromal populations (Fig. 4D, S6G-I). Compositional analysis revealed a significant shift in the type 2 diabetic islet vascular niche, characterized by expansion of islet-associated fibroblasts and a corresponding reduction in pericytes. These changes were significant at the donor level, whereas the composition of the exocrine vascular compartment remained largely unchanged (Fig. 4E, S6J). To further assess changes within the endocrine niche, we directly compared islet-associated fibroblasts and pericytes and observed a significant increase in the islet fibroblast-to-pericyte ratio in type 2 diabetic islets relative to non-diabetic controls (Fig. 4F). In contrast, the endothelial-to-pericyte ratio, the combined vascular-to-endocrine cell ratio, and the relative contribution of pericytes to the total vascular compartment were not significantly altered between non-diabetic and type 2 diabetic islets (Fig. S6K).

To better define vascular niche remodeling at the gene program level, we curated five transcriptional modules from genes differentially expressed between type 2 diabetic and non-diabetic vascular-associated populations in the PancDB dataset. To enable direct evaluation in the spatial dataset, only genes represented within the MERFISH panel were included and were assigned to modules based on Reactome and Gene Ontology annotations[100, 101], and established markers of pericytes, fibroblasts, and ECM programs[19, 25]. This process generated five transcriptional modules: Contractile, ECM Remodeling, Vascular BM, Interstitial ECM, and Cell Stress/Inflammatory (Fig. 4G). Within type 2 diabetic islet pericytes, stress/inflammatory and contractile program scores were increased relative to non-diabetic controls, whereas vascular BM, interstitial ECM, and ECM-remodeling module scores were reduced (Fig. 4H). Exocrine pericytes similarly exhibited increased contractile program scores together with reduced vascular BM, interstitial ECM, and ECM-remodeling signatures. In contrast to islet pericytes, however, stress/inflammatory signaling was decreased in type 2 diabetic exocrine pericytes relative to non-diabetic controls. We next examined islet-associated fibroblasts in type 2 diabetes and observed broad reductions across most transcriptional modules, accompanied by increased contractile and stress/inflammatory program scores, mirroring the pattern observed in islet pericytes. Exocrine fibroblasts displayed a similar overall profile but additionally showed increased ECM-remodeling and interstitial ECM program scores (Fig. 4I).

To validate the upregulation of interstitial collagens identified by spatial transcriptomics, we performed COL1A2 immunofluorescence staining in non-diabetic and type 2 diabetic pancreatic sections. Consistent with MERFISH analyses, COL1A2 fluorescence intensity within chromogranin A (CHGA)-defined islet regions was increased in type 2 diabetic islets compared with non-diabetic controls (Fig. 4J, S6L, Table S21). Together, these findings support coordinated remodeling of the islet vascular niche in type 2 diabetes through both ECM accumulation and altered vascular transcriptional states.

## Discussion

In this study, we have combined donor-resolved spatial transcriptomics with orthogonal validation using curated and re-integrated human pancreas reference datasets (PancDB[70] and Craig-Schapiro[25]) to define the cellular sources and spatial organization of ECM components within the adult human pancreas and to examine how these programs are further remodeled in type 2 diabetes. We found that pericytes, rather than ECs, exhibit the strongest BM-associated transcriptional program within the human islet vascular niche, including enrichment of *LAMA4* and *LAMB2*, laminin subunits previously reported to localize preferentially to the human intra-islet vascular BM[4]. ECs were distinguished by enrichment of *LAMA5*, a laminin subunit identified within both the peri-islet and the double intra-islet vascular BM[4]. In parallel, islet-associated fibroblasts exhibited enrichment of *LAMB1* and *LAMC1*, laminin proteins previously observed throughout the continuous peri-islet BM[1, 4]. The concordance between these cell type-specific transcriptional programs and prior protein localization studies supports distinct cellular contributions to human islet BM assembly. Together, these findings challenge the prevailing endothelial cell-centric model of islet BM production and instead support a model in which BM organization arises through coordinated contributions from multiple spatially specialized vascular- and stromal-associated populations.

The identification of a specialized islet-associated fibroblast population is particularly notable. Previous studies have proposed that distinct fibroblast-like populations may contribute separately to the peri-islet BM and the more collagen-rich intra-islet ECM, and our findings provide direct evidence supporting this concept[19, 102]. We show that islet-associated fibroblasts are enriched adjacent to the islet boundary and are transcriptionally distinct from broader exocrine fibroblast populations, suggesting specialized functions within the endocrine microenvironment. Their intermediate BM-associated transcriptional profile, marked by enrichment of *LAMC1* and *LAMB1*, ECM components preferentially associated with the peri-islet BM, suggests that islet-associated fibroblasts may contribute structurally to the peri-islet BM and potentially participate in the formation or maintenance of the characteristic double-layered BM architecture observed in human islets[1, 4]. Collectively, this further supports a model in which the endocrine ECM is maintained through coordinated contributions from multiple spatially specialized populations rather than a single dominant matrix-producing cell type.

Despite maintaining a conserved core BM-associated program, pericytes also demonstrated niche-dependent specialization between endocrine and exocrine compartments. Exocrine-associated pericytes displayed higher expression of genes linked to interstitial matrix remodeling and inflammatory signaling (*C1R*, *C1S*)[92], whereas endocrine-associated pericytes preferentially expressed genes associated with vascular regulation and endocrine support (*LAMC2*, *ACE2*)[95, 103]. These findings suggest that local microenvironmental cues may shape pericyte transcriptional identity within the pancreas, tuning ECM production and vascular-supportive functions according to regional tissue demands. These observations further support the growing view that pericytes do not represent a single homogeneous population but instead acquire niche-dependent phenotypes shaped by local vascular and metabolic environments[104–106].

To determine whether these vascular-supportive populations are altered in disease, we next examined remodeling in type 2 diabetes. In the droplet dataset, reduced *PDGFRB* expression in pericytes was particularly notable[70]. PDGFRB signaling is essential for pericyte survival, endothelial communication, and perivascular retention, and disruption of the PDGFB–PDGFRB axis is a recognized feature of diabetic microvascular disease across multiple tissues[96, 107–110]. Reduced PDGFRB expression, together with increased expression of *MYL9* and *MGP*, suggests a shift toward a dysfunctional or contractile pericyte state characterized by altered endothelial-pericyte interactions[99, 111, 112]. We additionally observed increased expression of *COL18A1*, a BM collagen that serves as the precursor of the anti-angiogenic fragment endostatin[113]. Elevated endostatin levels have been reported in diabetic microvascular complications of the cornea, retina, and kidney, where excessive anti-angiogenic signaling is thought to impair vascular adaptation and tissue repair[114–116]. Although our transcriptomic data do not directly assess endostatin production or activity, these findings suggest that diabetic pericyte dysfunction may extend beyond altered matrix deposition to include pathways regulating vascular maintenance and remodeling.

Spatial transcriptomics analyses further revealed coordinated remodeling of the endocrine vascular niche at both the compositional and transcriptional levels. Diabetic islets exhibited increased proportions of islet-associated fibroblasts together with an altered fibroblast-to-pericyte balance, suggesting selective restructuring of the endocrine stromal compartment. Diabetic islet pericytes exhibited reduced BM and ECM-associated programs alongside increased contractile, stress, and inflammatory signatures. Notably, these changes appeared specific to the endocrine niche, as exocrine pericytes also exhibited increased contractile programs but showed reduced stress and inflammatory signatures in type 2 diabetes, indicating that contractile remodeling may represent a shared adaptation, whereas inflammatory activation is selectively enriched within the endocrine microenvironment. Supporting this, spatial analysis showed that islet-associated fibroblasts and islet pericytes exhibited highly concordant transcriptional responses to type 2 diabetes, including reduced ECM programs and increased stress, inflammatory, and contractile signatures. Together, these findings indicate that type 2 diabetes remodels the endocrine vascular niche through both altered cellular composition and convergent shifts in vascular-supportive cell states.

Future work is needed to determine whether the transcriptional remodeling observed in type 2 diabetic pericytes directly contributes to beta cell dysfunction or vascular dysfunction or instead represents a secondary adaptation to the diabetic microenvironment. In particular, understanding whether restoration of canonical BM-associated pericyte programs can preserve endocrine niche integrity may have important therapeutic implications. The islet-associated fibroblast population identified here also warrants further investigation with respect to its developmental origins and potential role in peri-islet fibrosis[117]. In addition, the increased expression of contractile-associated genes observed in these cells raises the possibility that they may actively influence islet blood flow, vascular stability, and the mechanical properties of the endocrine niche in the diabetic pancreas[118]. Integrating higher-plex spatial transcriptomics, spatial proteomics, and functional perturbation studies will be important for determining how these phenotypic changes affect the vasculature and for resolving how distinct vascular-associated populations cooperatively regulate BM organization, ECM remodeling, and endocrine cell function in both healthy and diabetic human pancreata.

The findings reported here have direct implications for both disease modeling and regenerative medicine. Current efforts to generate vascularized islet organoids or engineer replacement tissues typically emphasize endothelial incorporation while largely overlooking the BM-producing role of fibroblasts and pericytes[119, 120]. Our data suggest that optimal reconstruction of a functional human endocrine vascular niche may require inclusion of cell-type-specific components. The identification of distinct ECM programs additionally provides a framework for designing biomaterials and engineered microenvironments that more faithfully reproduce the structural organization of the native human endocrine niche[121, 122].

### Limitations

Although our study includes human tissue from multiple donors, the number of human samples remains limited, reflecting the challenges of acquiring high-quality FFPE adult human pancreatic tissue with careful control over collection, timing, temperature, and fixation[123]. In addition, our spatial analyses rely on a targeted MERSCOPE gene panel rather than whole-transcriptome profiling, although recent work has shown this may be advantageous over broader targeted panels, as smaller gene panels offer greater sensitivity and can resolve major cell types[124]. ECM genes present additional technical challenges in spatial transcriptomics. Many ECM transcripts are large, structurally complex, and expressed at relatively low per-cell copy number, which can reduce detection sensitivity in FFPE tissue[125]. Furthermore, segmentation-based transcript assignment near vascular boundaries may imperfectly resolve closely opposed endothelial and pericyte transcripts, particularly in regions of dense BM deposition. Performing comparisons of our data with PancDB data, together with our donor-level pseudo-bulk analyses, only partially mitigates these concerns. Additionally, RNA abundance does not directly measure matrix deposition, crosslinking, or mechanical properties of the BM. However, confidence in our spatial findings is strengthened by cross-validation of all differentially expressed genes in at least one independent droplet-based single-cell sequencing dataset [25, 70].

Finally, while transcriptional enrichment of BM, ECM, and contractile gene programs suggests functional remodeling in type 2 diabetes, gene expression alone does not directly quantify contractile force generation, vascular perfusion dynamics, or ECM protein deposition. Future studies incorporating functional vascular assays, higher-depth spatial protein profiling, and longitudinal sampling will be important to further define the physiological consequences of the remodeling programs identified here[126].

## Supporting information

ESM

## Acknowledgments

We are grateful to the organ donors and their families whose contributions made this research possible. We thank Gregory Szot and the University of California, San Francisco Diabetes Research Center Islet Production Core Facility (RRID:SCR_015106) for providing human pancreatic tissue samples and the Gladstone Histology and Light Microscopy Core (RRID:SCR_017940) for assistance with tissue processing, sectioning, and staining. We also acknowledge the investigators and participants who generated and made publicly available the reference datasets used in this work, including PancDB, the Human Pancreas Analysis Program (HPAP; RRID:SCR_016202), the Craig-Schapiro human pancreas atlas, CellxGene (RRID:SCR_021059), and other publicly available human pancreas transcriptomic resources.

## Funding

This work was supported by grants funded by the Chan Zuckerberg Investigator Program to Z.J.G., the UCSF Center for Cellular Construction (DBI-1548297) to Z.J.G., the NIH/NIDDK (U01 DK135019-01) to J.B.S. and Z.J.G., and from Breakthrough T1D (Northern California Breakthrough T1D Center of Excellence; grant 4-COE-2024-1577-2-B) to J.B.S. We also acknowledge funding for the UCSF Diabetes Research Center Islet Core from NIH/NIDDK grant P30 DK135103. L.K.M. was supported by a National Science Foundation (NSF) Graduate Research Fellowship and the UCSF California Institute for Regenerative Medicine (CIRM) pre-doctoral fellowship (Grant EDUC4-12812).

## Data Availability

The MERFISH spatial transcriptomics dataset generated in this study has been deposited in the Gene Expression Omnibus (GEO) under accession number GSE333737 and is publicly available. Processed AnnData objects, cell-type annotations, custom MERFISH gene panel information, and analysis notebooks used for data processing, visualization, differential expression analysis, spatial analyses, and figure generation are available through the GEO submission and a public GitHub repository (https://github.com/lkim29meneses/Merscope_HumanPancreas_NBs). Publicly available reference datasets used in this study, including PancDB, HPAP, and the Craig-Schapiro human pancreas atlas, are available through their original publications and associated repositories. Additional data supporting the findings of this study are available from the corresponding authors upon reasonable request.

## Author Contributions

L.K.M., Z.J.G., and J.B.S. conceived and designed the study. L.K.M. performed all MERSCOPE experiments. H.J.K. and L.K.M. developed the computational pipelines and Jupyter notebooks used for MERSCOPE data analysis. G.L.S. provided human pancreatic tissue through the Human Islet Core at UCSF. L.K.M., Z.J.G., and J.B.S. wrote the manuscript. All authors reviewed, edited, and approved the final version of the manuscript.

## Ethics Statement

All human pancreatic tissue samples were obtained from de-identified donors following informed consent from the donor or, in post-mortem cases, the donor’s next of kin. Tissue procurement and handling were conducted under Institutional Review Board (IRB) or Independent Ethics Committee (IEC) approval in accordance with applicable ethical and regulatory guidelines. Donor information was fully anonymized prior to receipt by investigators.

## Study Design

Comparisons were performed between non-diabetic (n = 7) and type 2 diabetic (n = 5) pancreatic donor tissue samples. Additional analyses compared endocrine (islet) and exocrine compartments and vascular-associated cell populations, including endothelial cells, pericytes, fibroblasts, and islet-associated fibroblasts. Non-diabetic donors served as the control group. The primary experimental unit was an individual donor. For spatial transcriptomics analyses, cells were nested within donors, and donor-level replication was used for statistical comparisons where applicable (e.g., disease-group comparisons). Sample size was determined by the availability of suitable human pancreatic donor tissue and practical considerations associated with high-resolution spatial transcriptomics profiling. Formal power calculation was not performed. Donors were selected based on tissue availability, disease status, and RNA quality. Cell-level quality control filtering followed predefined criteria described in the Methods section. No donors meeting inclusion criteria were excluded. The exact n for each analysis is reported in the corresponding figure legends. Investigators were aware of donor disease status during sample processing and analysis, as blinding was not feasible given the use of predetermined human donor tissue. All cell type annotation, spatial, and statistical analyses were performed using standardized computational pipelines applied uniformly across samples to minimize analyst-introduced bias.

## Notes

https://www.ncbi.nlm.nih.gov/geo/query/acc.cgi?acc=GSE333737

### Competing Interest Statement

The authors have declared no competing interest.

