## Supplementary material for "The Spatial Landscape of Extracellular Matrix Gene Expression in Healthy and Type 2 Diabetic Human Pancreas": ESM

#### Supplementary Results

##### Quality Control, Integration, and Identification of Mesenchymal Populations in Human Pancreas Reference Datasets

To establish a reference framework for vascular-associated stromal populations, we first analyzed the PancDB [1]human pancreas scRNA-seq atlas. Following doublet removal and donor-level quality control, 46,106 cells were retained for downstream analysis (Fig. S1A-D). Examination of BM gene expression across major pancreatic cell types confirmed that endocrine populations exhibited minimal BM-related transcriptional programs, whereas mesenchymal and endothelial populations showed the highest expression of BM-associated genes. Although ductal cells displayed modest BM gene scores, this signal was largely attributable to *COL18A1* rather than coordinated expression of a broader BM transcriptional program (Fig. S1E).

Subsetting the mesenchymal compartment identified two major stromal lineages, designated MSL1 and MSL2 (Fig. S1F-J). MSL1 exhibited enrichment of canonical fibroblast-associated genes including *DCN*, *LUM*, *COL1A1*, and *COL1A2*, whereas MSL2 was characterized by expression of mural cell markers including *RGS5*, *CSPG4*, and *MCAM*. Cross-referencing with external human pancreas datasets demonstrated consistent transcriptional relationships between fibroblast, stellate, smooth muscle, and pericyte populations across studies (Fig. S1K, L).

##### Spatial Identification of Vascular-Associated Stromal Populations

To define the cellular sources of vascular ECM within the human pancreas, we generated a custom 300-gene MERSCOPE panel enriched for extracellular matrix, vascular, endocrine, and stromal-associated genes (Fig. S2A)[2–6]. Following integration of seven non-diabetic (ND) adult human donor datasets using CONCORD, major pancreatic cell populations segregated according to expected transcriptional identities while exhibiting minimal donor-driven clustering after using CONCORD instead of scVI (Fig. S2B-E).

Focused analysis of vascular-associated populations identified transcriptionally distinct endothelial cells, pericytes, fibroblasts, and an islet-associated fibroblast (IAF) population (Fig. S2F-I). IAFs expressed high levels of fibrillar collagen genes, basement membrane components, and ECM-remodeling factors, including *COL1A1*, *COL1A2*, *COL6A3*, *LAMA2*, *LAMC3*, *PRELP*, and *FBLN1*. Spatial analyses demonstrated enrichment of IAFs near the islet boundary, where they localized near vascular structures but remained spatially distinct from ECs and pericytes (Fig. S2J-N). These findings support the localization of IAFs within the peri-islet vascular niche, with 67.5%

located within 5 μm of the islet boundary, and are consistent with their proposed role in maintaining the islet vascular BM.

#### Cross-Dataset Validation of Islet-Associated Fibroblasts

To determine whether the IAF population represented a reproducible biological cell state rather than a spatial transcriptomic artifact, we interrogated the PancDB reference dataset. Re-clustering of fibroblast populations identified multiple transcriptionally distinct fibroblast subpopulations, including ECM-rich clusters characterized by elevated expression of *COL1A1*, *COL1A2, COL6A3, LAMB1, LAMC1, LAMC3, LUM, PRELP*, and *FBLN1* (Fig. S3A-D). Consolidation of these populations identified a fibroblast subset with a transcriptional profile closely resembling the spatially defined IAF population (Fig. S3E, F).

We next validated this population using the anatomically annotated Craig-Shapiro[2]human pancreas dataset. Islet fibroblasts displayed increased expression of ECM-associated genes and demonstrated enrichment of the candidate IAF signature identified in the spatial dataset (Fig. S3G). Comparison of average gene expression profiles between spatial IAFs and Craig-Shapiro islet fibroblasts revealed strong concordance across shared genes (Pearson $r = 0.738$; Spearman $\rho = 0.684$), providing independent support for a transcriptionally distinct fibroblast population associated with the endocrine niche (Fig. S3H).

#### Compartment-Specific ECM Programs in Islet Vascular Populations

To further define ECM specialization within the endocrine niche, we compared endothelial cells, pericytes, and IAFs. Endothelial cells preferentially expressed canonical vascular genes including *PLVAP, PECAM1, VWF, FLT1, ENG*, and *CD93*, whereas pericytes showed enrichment of ECM-associated genes including *PDGFRB, C1R*, *C1S, ACE2*, and *MFGE8* (Fig. S4A, B). In contrast, IAFs exhibited the strongest enrichment of fibrillar collagen programs, including *COL1A1, COL6A3,* and *FN1* (Fig. S4C).

Comparison of islet and exocrine pericytes identified compartment-specific transcriptional programs consistent with microenvironment-dependent specialization (Fig. S4D), although gene set scoring demonstrated no significant differences in the proportion of BM-high and ECM-high pericytes between endocrine and exocrine compartments (Fig. S4E).

#### Independent Validation of Vascular Cell States and ECM Programs

Analysis of the Craig-Shapiro dataset independently recapitulated the transcriptional distinction between endothelial cells, fibroblasts, and pericytes observed in the spatial data (Fig. S5A-D). ECs were enriched for vascular-associated genes, whereas pericytes retained canonical mural cell markers including *RGS5, TINAGL1, FLT1, COL6A1*, and *COL6A2*. Fibroblasts exhibited increased expression of fibrillar collagen and ECM-associated genes, including *COL1A1*, *COL1A2*, and *LAMA2* (Fig. S5D).

Within the pericyte population, islet- and exocrine-associated pericytes exhibited distinct transcriptional programs (Fig. S5E, F). Islet-associated pericytes showed increased expression of vascular-associated genes including *ACE2, RGS5, MFGE8*, and *LAMA2*, whereas exocrine pericytes displayed enrichment of ECM-remodeling and complement-associated genes including *IGFBP2, C1R, C1S, COL12A1, PCOLCE*.

**Mesenchymal Cells in Type 2 Diabetes Show Remodeled ECM Programs but Preserved Islet Vascular Architecture**

To investigate diabetes-associated changes in vascular ECM organization, we compared non-diabetic and T2D mesenchymal populations in PancDB. Differential expression and pathway enrichment analyses demonstrated activation of ECM-remodeling pathways in type 2 diabetes, including collagen formation, collagen fibril organization, ECM proteoglycans, TGFβ signaling, and epithelial-mesenchymal transition pathways (Fig. S6A-F). These changes were accompanied by distinct transcriptional remodeling of fibroblast- and pericyte-associated programs.

Spatial analysis of T2D samples identified transcriptional alterations across vascular-associated populations while preserving overall vascular organization (Fig. S6G-I). Quantification of endothelial-to-pericyte ratios, vascular coverage, pericyte abundance, and pericyte positioning relative to endothelial cells revealed no significant differences between non-diabetic and type 2 diabetic islets (Fig. S6J). Additional representative IF images further demonstrated increased intra-islet COL1A2 staining in type 2 diabetic islets compared with non-diabetic controls (Fig. S6K), supporting the increased COL1A2 expression observed by spatial transcriptomic analysis.

#### Supplementary Figure Legends

**Supplementary Figure S1. Quality control, integration, and identification of mesenchymal populations in the human pancreas scRNA-seq atlas.**

**(A)** Initial CONCORD-integrated UMAP embedding of the full PancDB pancreas dataset colored by major cell type annotations (top), donor identity (middle), disease state (middle), age (middle), and doublet metrics (bottom panels). Filtering thresholds for total counts and doublet score are indicated in red circles. Histograms show distribution of doublet scores across the dataset.

**(B)** CONCORD integrated PancDB atlas, prior to donor filtering but after removal of doublets and low-quality cells. UMAP embeddings are colored by cell type (top) or donor identity (middle). Donor quality control metrics include total cell number per donor, QC score distribution per donor (see Methods), and n_genes_by_counts (number of genes detected per cell) per donor.

**(C)** Sequential donor filtering strategy used for subset quality control. UMAP embeddings show progressive removal of low-quality donors (HPAP93, HPAP67, and HPAP27), resulting in the final filtered PancDB dataset used for downstream analyses in Figure 1 (n = 46,106 cells).

**(D)** Final integrated UMAP embedding following donor filtering, colored by annotated cell types (top), donor identity (middle), and doublet score metrics (bottom).

**(E)** Violin plot showing expression of top BM-associated genes within ductal epithelial cells, demonstrating selective enrichment of *COL18A1* relative to other canonical BM genes.

**(F)** UMAP embeddings of the final mesenchymal subset colored by total counts, number of detected genes, and doublet score.

**(G)** Leiden clustering of mesenchymal populations identifying eight stromal subclusters (n = 2,763 cells) within the integrated mesenchymal compartment.

**(H)** Top genes expressed in the mesenchymal subclusters used to annotate the populations into MSL1 and MSL2.

**(I)** UMAP embedding showing classification of mesenchymal populations into MSL1 and MSL2 stromal states by projection of canonical pericyte and fibroblasts markers.

**(J)** Heatmaps showing differential gene expression patterns across MSL1 and MSL2 populations. Expression values are scaled independently for each gene across populations.

**(K)** Table summarizing the external human pancreas reference datasets used for cross-dataset validation, including the PancDB, Human Cell Atlas (HCA), Tabula Sapiens, and Craig-Shapiro datasets.

**(L)** Venn diagram showing overlap of marker gene expression among fibroblast, stellate, smooth muscle, and pericyte populations across external pancreas reference datasets. Colored labels indicate representative genes enriched within activated or quiescent populations.

**Supplementary Figure S2. Spatial transcriptomic identification and characterization of vascular-associated stromal populations in the human pancreas.**

**(A)** Composition of the custom MERSCOPE spatial transcriptomics panel showing the relative distribution of ECM-associated genes, endothelial markers, endocrine markers, and additional lineage-specific genes included in the 300-gene panel design.

**(B)** Representative quality control metrics across seven donor spatial datasets. Left panels show segmented cell maps for each donor sample. Histograms display distributions of total transcripts per cell, unique transcripts per cell, transcripts per field of view (FOV), and segmented cell volume across samples.

**(C)** Representative MERSCOPE DAPI image of a pancreatic islet (Figure 2A) with Cellpose 2.0 segmentation-derived cell masks overlaid (thin colored outline) and colored according to assigned cell identity

(**D**) Comparison of donor integration using scVI and CONCORD. Left, scVI embedding colored by donor identity. Right, CONCORD-integrated embedding demonstrating improved donor mixing across shared cell populations.

**(E)** UMAP embedding of integrated spatial transcriptomic data colored by major pancreatic cell populations. Feature plots show expression of representative endocrine, exocrine, endothelial, mesenchymal, and immune marker genes across the embedding.

**(F)** Dot plot showing expression of curated marker genes across annotated pancreatic cell populations. Dot size represents fraction of cells expressing each gene and color indicates mean expression level.

**(G)** (*top*) UMAP visualization of healthy vascular cells colored by clusters after Leiden clustering colored by Leiden clustering. (*bottom*) Dot plot showing expression of representative marker genes across vascular-associated subclusters. Dot size indicates the fraction of expressing cells and color indicates mean expression level. Grey box highlights exocrine fibroblast populations, while the black box and asterisks denote islet-associated fibroblast (IAF) population.

**(H)** UMAP embedding of vascular-associated populations colored by anatomical location and cell type. Feature plots show expression of representative vascular and stromal marker genes, including *CHGA*, *PECAM1*, *JUN*, *RGS5*, *COL1A1*, *LUM*, *DCN*, *CSPG4*, *COL6A3*.

**(I)** Top markers expressed by spatial IAF cells in compassion to other vascular-associated populations.

**(J)** Dot plot showing expression of representative vascular, stromal, and BM/endocrine associated marker genes across endothelial cells, fibroblasts, IAF, and pericytes. Dot size represents the fraction of cells expressing each gene and color indicates mean expression level within each population.

**(K)** Histogram showing the distribution of shortest edge-to-edge distances between IAF and the nearest islet boundary. Negative distances indicate cells located within the islet compartment, whereas positive distances indicate cells located outside the islet boundary. The dashed red line denotes the islet boundary (0 μm). IAF were strongly enriched adjacent to the islet boundary, with the highest density observed near the endocrine-exocrine interface.

**(L)** Quantification of vascular-associated cell enrichment within ±5 μm of the islet boundary at the donor-paired level. Boxplots show the percentage of endothelial cells, pericytes, exocrine-associated fibroblasts, and IAF localized within 5 μm of the islet boundary across donors. Each point represents one donor.

**(M)** Quantification of distances from vascular-associated cell populations to the nearest EC within islets at the cell level. Pericytes localize significantly closer to ECs compared to islet-associated fibroblasts. Statistical significance was assessed using a Mann-Whitney U test.

**(N)** Quantification of islet endothelial coverage by pericytes within the islet vascular niche. Boxplot shows the fraction of endothelial cells located within 5 μm of a pericyte across donors. Each point represents one donor.

**(O)** Representative spatial map of vascular-associated populations surrounding a human islet. Colors indicate annotated pericytes (magenta), endothelial cells (green), fibroblasts (cyan), and ductal cells (yellow). Scale bar = 500 μm.

**Supplementary Figure S3. Cross-dataset validation of islet-associated fibroblasts using independent human pancreas scRNA-seq reference datasets.**

**(A)** Top differentially expressed genes defining the spatially identified islet-associated fibroblast (IAF) population from the MERSCOPE spatial transcriptomic dataset.

**(B)** Re-clustering of fibroblasts (MSL1) populations from the PancDB human pancreas scRNA-seq atlas. (*left*) UMAP showing the combined fibroblast compartment. (*right*) Leiden clustering identifies transcriptionally distinct fibroblast subpopulations.

**(C)** Differential expression analysis of individual PancDB fibroblast clusters. Clusters 2 and 5 were enriched for ECM- and basement membrane-associated genes including *COL1A1*, *COL1A2*, *COL3A1*, *COL4A1*, *COL5A1*, *SPARC*, *SERF2*, and *LGALS1*, whereas cluster 3 was enriched for canonical pericyte markers including *RGS5*, *IGFBP7*, and *SPARCL1*.

**(D)** (*left*) UMAP visualization of consolidation of PancDB fibroblast clusters into three major populations: fibroblasts, islet-associated fibroblasts, and pericytes. (*right*) Marker gene dot plot demonstrates that IAFs exhibit elevated expression of *COL1A1*, *COL1A2*, *COL6A3*, *LAMA2*, *LAMB1*, *LAMC1*, *LAMC3*, *PRELP*, and *GCG* relative to other fibroblast populations.

**(E)** Differential expression analysis of consolidated PancDB fibroblasts populations. IAFs were distinguished by enrichment of fibrillar collagen genes, BM-associated genes, and ECM-remodeling factors, whereas pericytes retained canonical mural cell markers and vascular-associated programs.

**(F)** Validation of fibroblast heterogeneity using the anatomically annotated Craig-Shapiro human pancreas scRNA-seq dataset. UMAPs colored by anatomical location (islet versus exocrine) and fibroblast identity. Differential expression analysis between islet and exocrine fibroblasts identified enrichment of endocrine and ECM-associated genes in islet fibroblasts. A candidate IAF signature derived from the spatial dataset, including *COL1A1*, *COL1A2*, *COL3A1*, *COL4A1*, *COL5A1*, *SPARC*, *SERF2*, and *LGALS1*, was similarly enriched in Craig-Shapiro islet fibroblasts.

**(G)** Correlation analysis comparing average gene expression profiles between spatially defined IAFs and Craig-Shapiro islet fibroblasts. Strong Pearson ($r = 0.608$) and Spearman ($\rho = 0.664$) correlations support the transcriptional concordance of these populations and provide independent validation of a distinct endocrine niche-associated fibroblast state.

**Supplementary Figure S4. Differential ECM and basement membrane gene expression across endothelial cells, pericytes, and islet-associated fibroblasts.**

**(A)** Representative MERSCOPE spatial transcriptomic image showing localization of BM genes surrounding islet vascular structures. Expression of *COL4A1*, *COL4A2*, and *LAMC1* is enriched in vascular-associated stromal cells adjacent to endocrine tissue.

**(B)** Differential expression analysis comparing islet endothelial cells and islet pericytes. Volcano plot highlights endothelial-enriched genes including *PLVAP*, *PECAM1*, *VWF*, *FLT1*, *ENG*, and *CD93*, while pericytes show enrichment of ECM- and BM-associated genes including *PDGFRB*, *COL6A1*, *COL5A3*, *C1R, C1S*, *ACE2*, and *MFGE8*.

**(C)** Differential expression volcano plot analysis comparing islet-associated fibroblasts and islet pericytes. Islet-associated fibroblasts were enriched for ECM genes including *COL1A1*, *COL6A3*, *COL16A1*, *MMP2*, and *LAMA2* whereas pericytes were enriched for pericyte and EC-associated genes including *CSPG4*, *ACE2*, *PLVAP*, *VWF*, and *CD93*.

**(D)** Differential expression analysis comparing islet and exocrine pericytes. Volcano plot demonstrates compartment-specific transcriptional programs, including enrichment of ECM and vascular-associated genes between endocrine (red)- and exocrine (purple)-associated pericyte populations.

**(E)** Quantification of BM and ECM gene set scores in islet and exocrine pericytes. The percentage of cells classified as BM-high or ECM-high was calculated using gene set scoring and threshold-based classification. P values (BM = 0.109; ECM = 0.578) were calculated using a paired Wilcoxon signed-rank test; ns, not significant.

**Supplementary Figure S5. Cross-platform validation of vascular and stromal populations using the Craig-Shapiro human pancreas scRNA-seq dataset.**

**(A)** UMAP visualization of the Craig-Shapiro dataset colored by cell originating location (islet or exocrine), cell type, and representative vascular, stromal, and endocrine (*GCG*) marker genes. Expression patterns identify fibroblast, endothelial, and pericyte populations and demonstrate conservation of transcriptional programs across datasets.

**(B)** (*top*) UMAP projections colored by anatomical location (islet versus exocrine) and annotated cell type. Islet and exocrine stromal populations form partially distinct transcriptional states within the fibroblast and pericyte compartments. (*bottom*) Gene expression of *LUM* and *RGS5* further show differential expression between the populations.

**(C)** Differential expression analysis between endothelial cells and pericytes within islet-associated populations. Volcano plot and corresponding heatmap demonstrate enrichment of endothelial markers (*CECAM1*, *HSPG2*, *SOX18*, *LAMA5*) in ECs and mural cell markers (*COL5A3*, *COL3A1*, *IRX2*, *COL1A2*, and *MFGE8*) in pericytes.

**(D)** Differential expression analysis between fibroblasts and pericytes within islet-associated populations. Pericytes preferentially express canonical mural cell genes including *RGS5*, *TINAGL1*, *FLT1*, and *TCIM*, whereas fibroblasts show increased expression of fibrillar collagen and ECM genes including *COL1A1*, *COL1A2*, and *LAMA2*. Heatmap summarizes representative marker genes.

**(E)** Analysis of pericyte heterogeneity in the Craig-Shapiro dataset. UMAP projections identify islet- and exocrine-associated pericytes. Differential expression analysis demonstrates compartment-specific transcriptional programs between endocrine (red)- and exocrine (purple)-associated pericytes.

**(F) (*left*)** Divergent bar plots showing the top ten differentially expressed genes between endocrine- and exocrine-associated pericytes. (*right*) Divergent bar plots of the same genes shown in Fig. 3H showing similar gene expression patterns as the MERSCOPE differential gene expression analysis between islet and exocrine-associated pericyte populations.

**Supplementary Figure S6. Type 2 diabetes-associated remodeling of vascular populations in the human islet.**

**(A)** Integration and quality control of vascular- and mesenchymal-associated populations from the PancDB human pancreas scRNA-seq atlas. UMAP projections are shown before and after donor-level quality control filtering. Cells are colored by donor identity, disease status, and major cell type annotation. Following quality filtering, 93,746 cells were retained for downstream analyses. Mesenchymal populations were subsequently extracted for focused analysis of diabetes-associated stromal remodeling.

**(B)** UMAP of the PancDB mesenchymal subset, highlighting fibroblast- and pericyte-associated populations used for downstream clustering and differential expression analyses.

**(C)** Expression of representative marker genes distinguishing fibroblasts and pericyte subpopulations. Feature plots show enrichment of ECM-associated genes (*COL6A3, COL1A2, DCN, LUM, ADAMTS4*), mural cell markers (*RGS5, TFPI*), and fibroblast-associated genes (*C11orf96, FABP4*), supporting separation of two major mesenchymal lineages (MSL1 and MSL2).

**(D)** Gene set enrichment analysis comparing PancDB T2D and non-diabetic fibroblasts and pericyte populations. Differentially expressed genes were ranked by log fold change and analyzed using curated Reactome, GO, and ECM-associated pathways. T2D-associated populations exhibited enrichment of collagen organization, ECM proteoglycan, collagen fibril assembly, and TGFβ-related pathways, consistent with matrix remodeling and fibrotic activation.

**(E)** Differential expression analyses comparing PancDB non-diabetic and T2D fibroblasts and pericyte populations. Volcano plots identify genes associated with ECM remodeling, basement membrane organization, vascular function, and inflammatory signaling that are altered in T2D.

**(F)** Heatmap of genes differentially regulated between fibroblasts and pericyte populations in PancDB non-diabetic and T2D samples. Distinct transcriptional programs associated with extracellular matrix organization, BM assembly, vascular remodeling, and inflammatory signaling distinguish diabetes-associated stromal states.

**(G)** UMAPs of spatial transcriptomic characterization of integrated MERSCOPE T2D pancreas-associated populations across five donors. UMAPs are colored by donor identity, anatomical location (endocrine-associated versus exocrine-associated), and major cell type annotation.

**(H)** Cell type composition gene expression dot plot across major pancreatic cell populations. Dot plot analysis demonstrates cell type-specific expression of representative endocrine, vascular, and immune genes.

**(I)** (*left*) UMAP of subset of integrated MERSCOPE vascular-associated populations across all 12 donors. (*right*) Dot plot of vascular-associated cell makers across the identified vascular-associated populations in the MERSCOPE dataset. Expression of canonical vascular markers (*PECAM1, PLVAP, VWF*) and mural/fibroblasts cell markers (*RGS5, CSPG4, DCN, LUM*) confirms population identities. (*bottom*) Heatmap of BM genes across T2D vascular-associated populations.

**(J)** Donor-level compositional analysis of vascular-associated cell populations in ND and T2D pancreas. Mean donor-normalized fractions of ECs, pericytes, and IAFs within islet and exocrine vascular compartments. Bars represent mean ± SD across donors, and individual points represent one donor. T2D islets exhibited a significant increase in IAF abundance and a corresponding decrease in pericyte abundance compared with ND islets (both $p = 0.002$). Statistical comparisons were performed using two-sided unpaired Mann-Whitney U tests.

(**K**) Quantification of vascular composition and spatial organization in ND and T2D islets. Metrics examined included endothelial-to-pericyte ratio, islet vascular coverage, pericyte fraction, and pericyte positioning relative to vascular structures. No significant differences were detected between ND and T2D samples for these structural vascular parameters. ns = not significant by two-sided unpaired Mann-Whitney U test.

**(L)** Further validation of increased intra-islet COL1A2 protein expression in type 2 diabetic human islets. Additional representative images of immunofluorescence staining for COL1A2 (magenta), CHGA (orange), COL4A1 (green), and DAPI (blue) in ND and T2D human pancreatic islets. Dashed lines indicate CHGA-defined islet boundaries used for quantification. Upper panels show representative T2D islets, which exhibit increased intra-islet COL1A2 staining relative to ND islets shown in the lower panels. Bottom rows show additional representative merged images from each condition. All images were acquired using identical staining, imaging, and acquisition settings. Scale bars = 40 μm.

**Supplementary Materials and Methods:**

**scRNA-seq Quality Control and Filtering:** scRNA-seq data (.h5ad) were downloaded from the PancDB website, and further filtering using Scanpy quality control was performed to exclude low-quality cells and technical artifacts. QC score was quantified using a composite score base total counts and number of detected genes; log10-transformed and z-score normalized. The final QC score was defined as: QC score = z (log(10(total_counts + 1)) + z (log10(n_genes_by_counts + 1)) Higher scores indicate cells with greater transcript detection and gene diversity (higher quality cells). Cells with ‘unknown’ cell label, fewer than 100 detected gene counts, or greater than 20,000 were removed to eliminate empty droplets and potential doublets, respectively. Doublets were identified using DoubletDetection. The expected number of doublets was estimated, and cells were classified as singlets or doublets using the doubletdetection () classification framework. Predicted doublets (score of 1) were excluded from further analysis.

**Refining PancDB dataset summary (Fig. S1):**

- Dataset was subsetted for adult control (ND) donors only - scviUMAP = 88,701 cells.
- Cells were filtered to those with >100 but < 20,000 counts; 75,763 cells.
- Cells labeled unknown were removed.
- Doubletdetection was run and doublets with a positive prediction score = 1 were removed; 71,639 cells.
- CONCORD integration.
- Filtering of low-quality samples; 46,106 cells, 22 donors.
- Mesenchymal cells totalled 3,010 cells.
- Before further analysis, we checked doublet _score since there were many small clusters of cells away from the main clusters and saw that those cells had relatively higher doublet scores; therefore, cells with a doublet score >20 were filtered out.
- Unsupervised clustering with resolution = 0.1 was then performed, which resulted in 8 clusters. Further examination showed that cluster 7& 8 were only 118 and 50 cells total, respectively.
- Before filtering out the small clusters, we analyzed DGE using sc.rank_genes and observed that cluster 7 was comprised mostly of ribosomal genes, while cluster 8 contained some fibroblast-like genes. This led us to remove cluster 7. Clusters 0, 1, 4, 5, 6, and 8 were merged and labeled as fibroblast-like MSL1, and clusters 2 and 3 were merged and labeled as pericyte-like MSL2.

**For T2D PancDB Analysis (Fig. S7):**

- The same quality control, filtering, doublet removal, and integration procedures described for the non-diabetic PancDB analysis were applied to the combined non-diabetic and T2D dataset. Samples HPAP109 and additional low-quality donors identified during quality control were excluded prior to downstream analyses.

- MSL1 (fibroblast-like) and MSL2 (pericyte-like) populations were identified using the same clustering strategy, marker gene criteria, and differential expression framework described for the non-diabetic dataset (Fig. 1).
- Subsequent analyses were performed to compare cell composition, gene expression programs, and pathway enrichment between non-diabetic and type 2 diabetic samples within each vascular-associated population.

**Gene Set Scoring:** Gene set scores were calculated using Scanpy's score_genes function. For each cell, a score was computed as the average expression of genes within the target gene set minus the average expression of expression-matched control genes. Scores were calculated using the expression matrix stored in adata.X (use_raw=False) and stored in adata.obs for downstream analyses. To ensure separation between matrix programs, genes contained within the BM gene set were excluded from the broader ECM gene set prior to scoring, such that ECM scores reflected non-BM matrix components. For visualization, mean gene set scores were scaled across cell types using Scanpy's standard_scale='var' option. In spatial analyses, BM genes are listed in Fig. 3A, and ECM genes used are listed in Table S2.

**MERFISH for FFPE tissue sample**s: A fully detailed, step-by-step guide to the MERFISH sample preparation full protocol is available at https://vizgen.com/resources/fresh-and-fixed-frozen-tissue-sample-preparation. Briefly, FFPE samples from selected donors were sectioned using a microtome (Leica Microtome, HM325) at 4-μm thickness and placed on a MERSCOPE FFPE slide (Vizgen, catalog no. 20400100) in accordance with Vizgen's MERSCOPE User Guide for FFPE samples. The tissue sections were then deparaffinized three times using Deparaffinization Buffer (Vizgen, catalog no. 20300112) at 55 °C for 5 min and washed three times with 100% ethanol. Each wash was for 2 min, followed by 2 min in 90% ethanol and a 2-min 70% ethanol rehydration step. The rehydrated tissue sections were incubated with Decrosslinking Buffer (Vizgen, catalog no. 20300115) at 90 °C for 15 min and cooled at room temperature for 10 min. The uncrosslinked tissue sections were then incubated with Conditioning Buffer (Vizgen, catalog no. 20300116) at 37 °C for 30 min, and Pre-Anchoring Reaction Buffer (Vizgen, catalog no. 20300113) for 2 h at 37 °C. Following anchoring pretreatment, tissue slices were stained for cell boundaries using Vizgen's Cell Boundary Kit (catalog no. 10400009) following the user guide. Samples were then washed with Formamide Wash Buffer (Vizgen, catalog no. 20300002) at 37 °C for 30 min and incubated with Anchoring Buffer (Vizgen, catalog no. 20300117) at 37 °C overnight. Following overnight incubation, samples were washed, then gel embedded and cleared. Tissues were cleared using the resistant-clearing protocol for the allowed maximum time (72 h).  After tissue clearing, samples were treated with MERSCOPE Photobleacher (Vizgen, catalog no. 10100003) for 4 h, washed with Formamide Wash Buffer at 37 °C for 30 min, and then incubated with MERSCOPE Gene Panel Mix for 48 h at 37 °C.

**Islet Annotations:** Manual islet annotation was performed in MERSCOPE Visualizer. Regions were selected using four criteria: (1) expression of endocrine-associated transcripts within the candidate region, (2) presence of DAPI signal confirming cellular localization, (3) presence of cells annotated as endocrine populations based on transferred cell labels displayed in the Visualizer overlay, and (4) a minimum of three endocrine cells per annotated islet. Regions meeting all criteria were annotated as islets. Annotated islet geometries and associated metadata, including transcript coordinates, cell identities, and islet assignments for each cell were exported as .csv files for downstream analyses (Tables S8 and S9). A total of 2,387 ND islets and 1,311 T2D islets were annotated.

**Clustering and Cell-type Annotation:** To construct integrated compartment-level datasets, islet-associated cells from all donors were concatenated to generate a combined islet dataset. Exocrine-associated cells from all donors were similarly concatenated to generate a combined exocrine dataset. Because the exocrine compartment contained substantially more cells than the islet compartment, exocrine cells were randomly downsampled (2.8%) to achieve comparable cell numbers between compartments and reduce potential clustering biases driven by unequal sampling. The balanced islet and exocrine datasets were subsequently combined into a single AnnData object for downstream integration and analysis. Spatial datasets were integrated using CONCORD with donor identity specified as the batch variable. A k-nearest neighbor graph was constructed using 15 neighbors and 30 principal components (n_neighbors = 15, n_pcs = 30). Two-dimensional embeddings were generated from the CONCORD latent space and graph-based clustering was performed using the Leiden algorithm (resolution = 1.0) to identify major cellular populations. Cluster identities were assigned using canonical marker genes and differential expression analysis performed with Scanpy's Wilcoxon rank-sum test implementation (sc.tl.rank_genes_groups).

- Endothelial and mesenchymal populations were subsetted from the integrated spatial transcriptomic dataset and re-clustered using Leiden clustering to resolve vascular-associated stromal heterogeneity. Initial clustering at a resolution of 0.5 did not adequately separate populations expressing distinct levels of the canonical pericyte marker *RGS5*. Therefore, cells were re-clustered at a higher resolution (resolution = 1.0), resulting in 14 transcriptionally distinct clusters.

- Cluster identities were assigned based on differential gene expression and canonical marker genes. Clusters were classified as ECs, pericytes, fibroblasts, or a distinct fibroblast population characterized by enrichment of ECM genes, BM-associated genes, and endocrine niche-associated transcriptional programs. This population remained transcriptionally distinct from both pericyte and exocrine fibroblast populations and was therefore annotated as islet-associated fibroblasts (IAFs).
- To evaluate anatomical localization, annotated cell populations were intersected with manually defined islet regions of interest (ROIs). Approximately 93% of IAFs were located within islet ROIs, whereas approximately 93% of exocrine fibroblasts were located outside of islet ROIs.

**Spatial Distance Calculations:** Spatial relationships between vascular-associated cell populations were quantified using segmentation-derived cell polygons and Shapely geometric operations. Cell boundaries generated from MERSCOPE segmentation masks were converted into polygon geometries and indexed using STRtree spatial indexing to enable efficient nearest-neighbor queries. Distances were calculated in microns using the native MERSCOPE coordinate system. For cell-to-cell analyses, the shortest edge-to-edge distance between cell boundaries was computed using polygon geometry. This approach measures the minimum physical separation between neighboring cells and avoids biases introduced by differences in cell size or morphology. Distances between ECs, pericytes, fibroblasts, and endocrine cells were calculated using nearest-neighbor searches across annotated cell populations. For islet boundary analyses, manually annotated islet polygons generated in MERSCOPE Visualizer were used to define endocrine compartment boundaries. The shortest edge-to-edge distance between each cell polygon and the nearest islet boundary polygon was calculated. Cells located within the annotated islet region were assigned negative distance values, whereas cells located outside the islet boundary were assigned positive values. Distances were summarized at both the cell level and donor level for downstream statistical analyses and visualization of vascular niche organization. To quantify vascular niche organization within pancreatic islets, nearest-neighbor analyses were performed between ECs, pericytes, and endocrine cells using segmentation-derived cell boundaries. Endothelial-pericyte interactions were assessed by calculating the shortest edge-to-edge distance between each endothelial cell and its nearest pericyte. Pericyte coverage was subsequently calculated as the proportion of ECs satisfying this criterion and summarized at the donor level.

**Immunofluorescence (IF) Staining of Human FFPE Pancreatic Tissue:** Formalin-fixed paraffin-embedded (FFPE) human pancreatic tissue sections (4-5 μm) were baked at 55°C for 15 minutes prior to processing. Sections were deparaffinized in xylene and rehydrated through graded ethanol washes to distilled water. Antigen retrieval was performed using heat-mediated retrieval in citrate buffer pH = 6.0 (Vector Labs, H-3300-250) for 15 minutes and then cooled to RT for another 15 minutes. Tissue sections were permeabilized and blocked in 0.5% PBST with 5% normal donkey serum (NDS) for 1 hour at RT. Primary antibodies were incubated overnight at 4°C in a humidified chamber. Following three 1X PBS washes, sections were incubated with species-specific AlexaFluor-conjugated secondary antibodies at RT for 1 hour. Nuclei were counterstained with DAPI if needed. Slides were mounted using antifade mounting media (Vector Labs, H-1000-10). Fluorescence imaging was performed using a Zeiss LSM800 laser-scanning confocal microscope. For colocalization analysis, images were processed in Fiji/ImageJ and quantified using the Coloc2 plugin. For COL1A2 analysis, islet regions were manually annotated in Fiji/ImageJ using CHGA endocrine marker staining as a guide. Mean COL1A2 fluorescence intensity was quantified within each manually defined region following background subtraction. Table of antibodies used is provided in SI (Table S22).

**RNAscope Multiplex Fluorescence Assay on Human FFPE Pancreatic Tissue:** RNAscope multiplexed fluorescence *in situ* hybridization was performed on FFPE human pancreatic tissue sections using the RNAscope Multiplex Fluorescent Reagent Kit (Advanced Cell Diagnostics, Cat. No. 323100 & 323270) according to the manufacturer's protocol[7]. Target-specific RNAscope probes were hybridized to tissue sections, followed by sequential signal amplification steps according to the manufacturer's instructions. Fluorescent detection was performed using Opal fluorophores (Akoya Biosciences). Where indicated, RNAscope was combined with immunofluorescence staining following the completion of probe amplification. Nuclei were counterstained with DAPI, and slides were mounted using an antifade mounting medium (Vector Labs, H-1000-10). Fluorescence imaging was performed using a Zeiss LSM800 laser-scanning confocal microscope at either 40x or 63x.

**Cell Composition Analysis:** Vascular-associated cell composition was assessed using annotated ECs, pericytes, fibroblasts, and islet-associated fibroblasts identified from the integrated spatial transcriptomic dataset. For each donor, the total number of cells belonging to each vascular-associated population was quantified and expressed as a proportion of all vascular-associated cells within the analyzed region. To account for differences in total cell recovery between donors, cell composition was calculated on a donor-by-donor basis prior to statistical testing. Endothelial-to-pericyte ratios were calculated by dividing the number of ECs by the number of pericytes identified within each donor. Islet vascularization was calculated by the ratio or ECs + pericytes to total endocrine cells from that sample. Similarly, the relative abundance of pericytes was calculated as the percentage of pericytes divided by total islet vascular-associated cells.

For disease-state comparisons, donor-level composition metrics were compared between ND and T2D samples using Mann-Whitney U tests.

**Gene set enrichment analysis:** We performed gene set enrichment analysis (GSEA) separately for MSL1 (fibroblast-like) and MSL2 (pericyte-like) populations using GSEApy. Pseudobulk differential expression was first computed by aggregating counts per donor within each mesenchymal subtype, followed by differential testing using a donor-level t-test on log-normalized CPM (counts per million) values. Genes were ranked by their pseudobulk t-statistic, which was chosen as the ranking metric over log-fold change alone to account for cross-donor variance. The ranked gene list was passed to gseapy.prerank with the following parameters: min_size=5, max_size=500, permutation_num=1000, and seed=1. Gene sets were drawn from MSigDB Hallmark (v.2023) and Reactome Pathways 2024 databases. Terms with a false discovery rate (FDR) q-value < 0.25 were considered significant, consistent with standard GSEA reporting thresholds. Normalized enrichment scores (NES) are reported with direction indicating enrichment in T2D (positive) or ND (negative). Results were visualized using matplotlib.

**Manual gene set curating for Type 2 Diabetes spatial analysis:** To characterize T2D-associated transcriptional remodeling in vascular-associated populations, custom gene programs were manually curated using differential expression results from the PancDB human pancreas single-cell RNA-seq reference dataset (Table S14-15). Differential expression analyses comparing type 2 diabetic and non-diabetic pericytes and fibroblasts were used to identify genes consistently associated with disease-related changes in ECM organization, BM maintenance, contractility, stress responses, inflammation, and matrix remodeling. Genes exhibiting significant or strong differential expression, together with genes present in the spatial gene panel and supported by prior literature, were grouped into biologically coherent transcriptional modules. Curated gene sets included BM, interstitial ECM, ECM remodeling, contractile, and stress/inflammatory programs. Gene program activity was quantified in spatial transcriptomic datasets using Scanpy's score_genes function and gene program scores were subsequently Z-score normalized across cells. These manually curated programs were used to compare healthy and type 2 diabetic tissues and to identify compartment-specific remodeling signatures within endocrine and exocrine vascular niches.

**Cross-Reference Validation with External scRNA-seq Datasets:** To validate spatially identified stromal and vascular-associated populations, comparisons were performed against external human pancreas scRNA-seq reference datasets, including PancDB and the Craig-Shapiro pancreas dataset. Spatial populations were compared against reference populations using shared genes present across datasets and restricted to genes included within the spatial transcriptomics panel where indicated. For transcriptional concordance analyses, average gene expression profiles were calculated for each annotated spatial and reference population. Pearson correlation and Spearman correlation metrics were used to quantify cross-platform similarity between matched populations. Gene-level comparisons were performed using normalized mean expression values across shared genes. Differentially expressed genes identified in spatial analyses were additionally cross-referenced against external reference datasets to reduce the likelihood that observed transcriptional differences reflected segmentation artifacts, transcript diffusion, or spatial contamination effects. Where PancDB lacked explicit anatomical annotations, corresponding stromal populations were identified computationally based on transcriptional similarity and marker gene enrichment. Spatially defined islet-associated fibroblasts were subsequently validated against anatomically annotated stromal populations in the Craig-Shapiro dataset.

- Cells previously annotated as MSL1 fibroblast-like populations in PancDB (n = 1,833 cells) were subsetted and re-clustered using Leiden clustering (resolution = 0.5) to further resolve fibroblast heterogeneity. This analysis identified six transcriptionally distinct clusters, each containing more than 100 cells.
- Differential expression analysis was performed for each cluster using the Wilcoxon rank-sum test. Two clusters (clusters 2 & 5) exhibited elevated expression of extracellular matrix and basement membrane-associated genes that closely resembled the spatially identified islet-associated fibroblast (IAF) population. Representative markers included *COL1A1*, *COL1A2*, *COL3A1*, *COL4A1*, *COL5A1*, *SPARC*, *SERF2*, *LGALS1*, *PRELP*, and additional matrix-associated genes. In contrast, cluster 3 displayed high expression of canonical pericyte markers including *RGS5*, *CSPG4*, *MCAM*, and *PDGFRB*.
- Based on these transcriptional profiles, clusters 2 & 5 were merged and annotated as IAF-like cells. Clusters 0, 1, & 4 were merged and annotated as fibroblasts, whereas cluster 3 was annotated as pericytes. Subsequent differential expression analyses demonstrated strong concordance between the PancDB IAF-like population and the spatially identified IAF population, including shared enrichment of fibrillar collagen genes, basement membrane-associated genes, and extracellular matrix remodeling programs. These findings provided independent validation of a transcriptionally distinct endocrine niche-associated fibroblast population.

Craig-Shapiro Dataset Analysis:

- The anatomically annotated human pancreas scRNA-seq dataset generated by Craig–Shapiro et al. was downloaded from the Gene Expression Omnibus (GEO) and processed in Scanpy. Major pancreatic cell populations were reconstructed using the published cell annotations. Endothelial cells, pericytes, and fibroblasts were subsequently subsetted and analyzed independently for cross-reference validation of spatially identified vascular-associated populations.

- To facilitate direct comparison with the MERFISH dataset, the reference dataset was restricted to genes represented in the custom 300-gene spatial transcriptomics panel. Gene expression values were normalized and log-transformed prior to downstream analyses.
- For validation of islet vascular populations, endothelial cells, pericytes, and fibroblasts were first restricted to cells annotated as originating from islet tissue. Expression of genes identified in the spatial dataset was assessed within the corresponding reference populations and visualized using violin plots and feature plots.
- To evaluate compartment-specific transcriptional programs, pericytes were further stratified by anatomical location (islet versus exocrine tissue). Differential expression analysis was performed using the Wilcoxon rank-sum test, and results were visualized using volcano plots. Genes identified as differentially expressed between islet and exocrine pericytes in the spatial dataset were subsequently examined in the Craig–Shapiro dataset to determine whether similar location-associated expression patterns were observed across platforms.

### Supplementary Figure 1

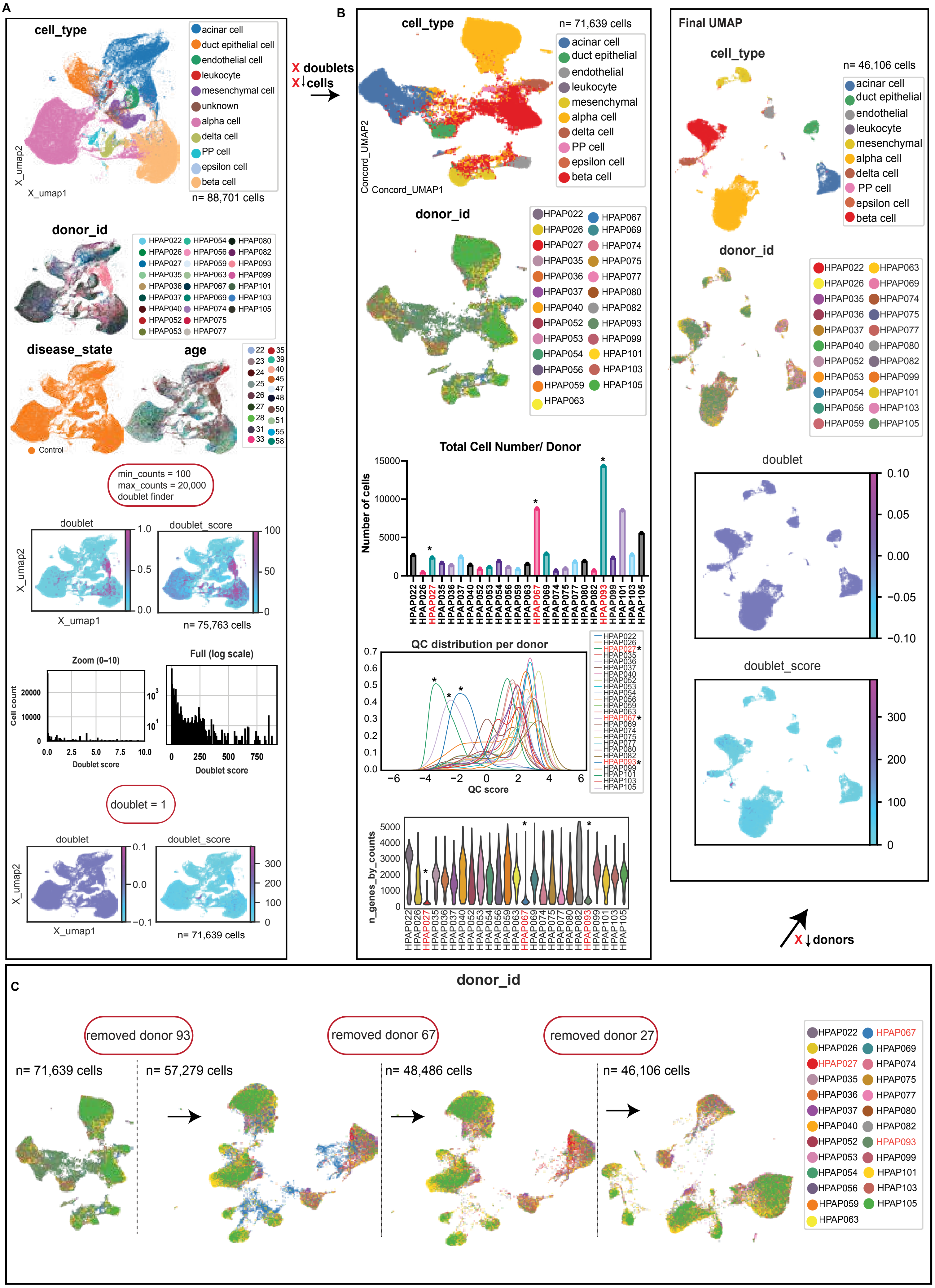

### Supplementary Figure 1

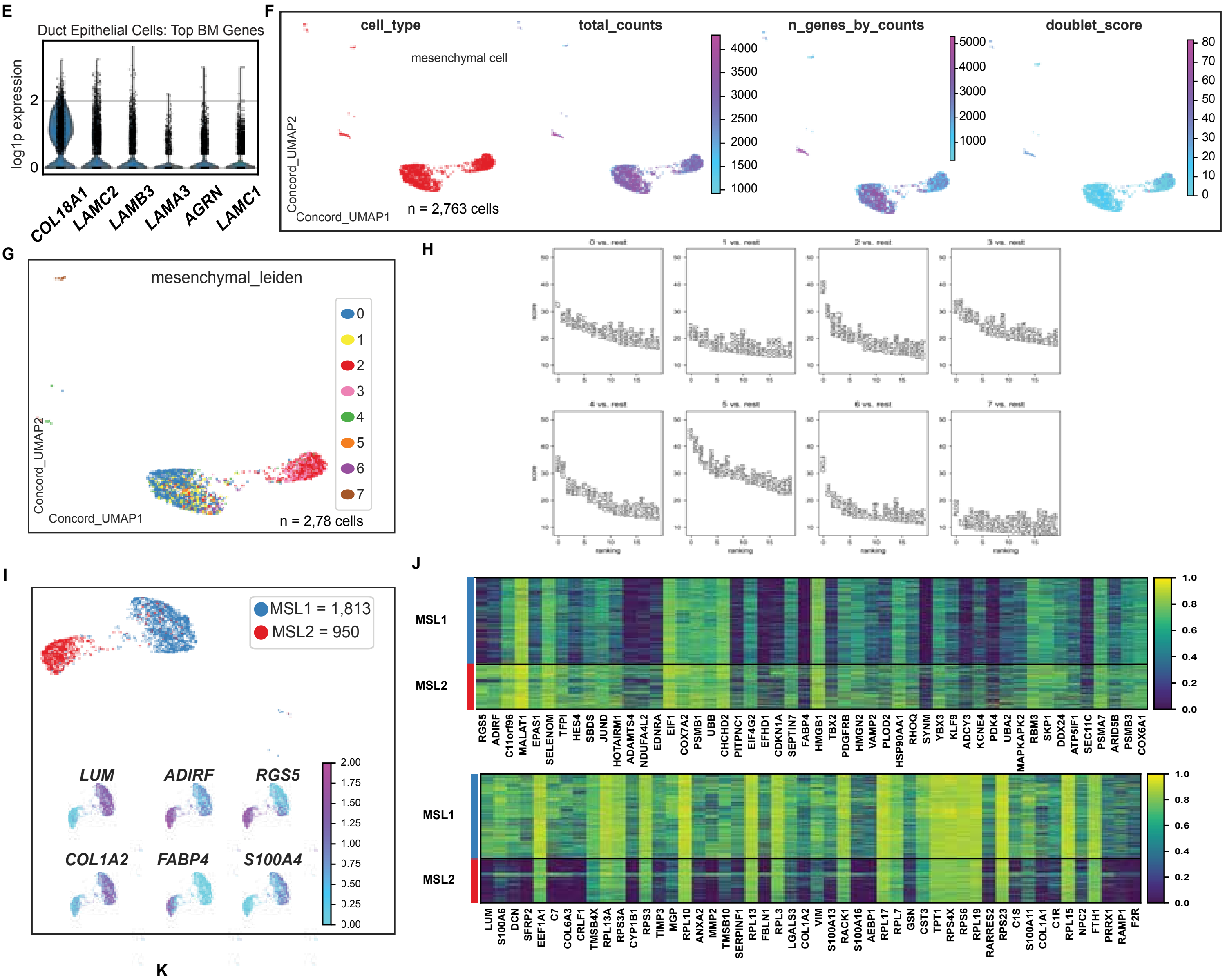


K

| Dataset | # of Cells | # of Genes | # of Donors | Subset | Link |
|---|---|---|---|---|---|
| PancDB | 2,523 | 21,481 | 23 | Yes | https://hpap.pmacs.upenn.edu/explore/download?matrix |
| HCA (Pancreas) | 2,208 | 33,053 | 16 | Yes | https://data.humancellatlas.org/hca-bio-networks/pancreas |
| Tabula Sapiens | 440 | 58,870 | 2 | Yes | https://figshare.com/articles/dataset/Tabula_Sapiens_v2/27921984 |
| Craig-Schapiro | 10,835 | 20,896 | 3 | Yes | https://www.ncbi.nlm.nih.gov/geo/query/acc.cgi?acc=GSE221237 |

L

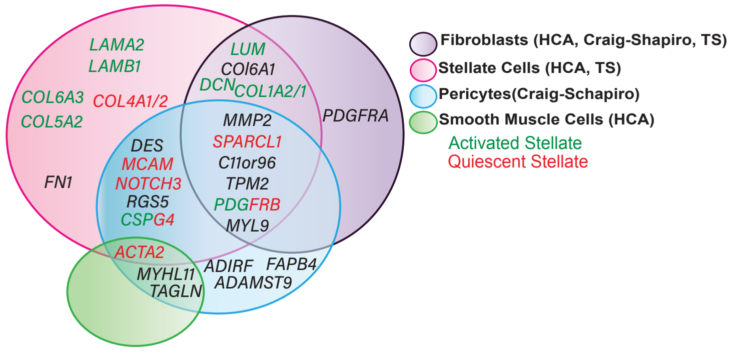

### Supplementary Figure 2

**A**

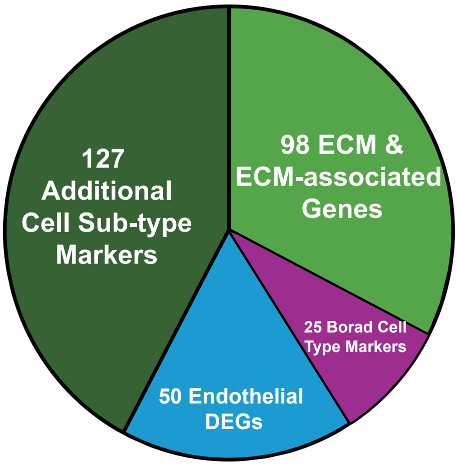


| 1. ECM Genes | 2. Canonical Cell Type Marker Genes | 3. Subpopulation Cell TypeGenes |
|---|---|---|
| Sufficinet Probe Binding Sites on RNA-sequence | Sufficinet Probe Binding Sites on RNA-sequence | Sufficinet Probe Binding Sites on RNA-sequence |
| FPKM Levels | FPKM Levels | FPKM Levels |
| Human Proteomics ECM | Human Cell Atlas | Human Cell Atlas |
| Human Protein Atlas | Cell x Gene | Cell x Gene |
| Mouse islet ECM RNA-seq | | Craig-Shapiro EC Atlas |

**B**

HuP-1 (HuP03A_L) = 644,325 cells

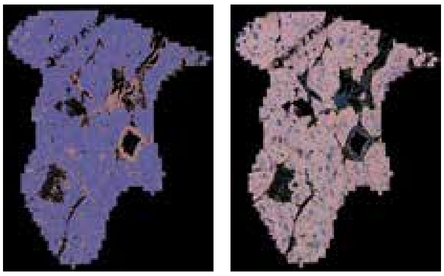

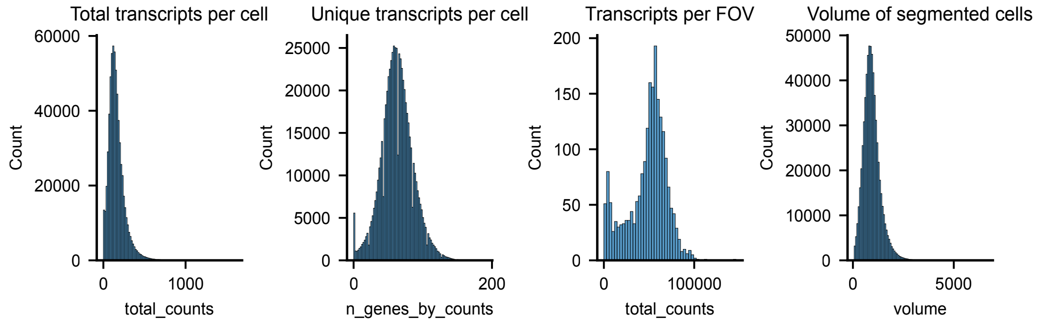


HuP-1 (HuP03A_S)

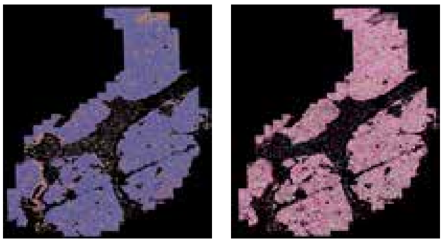

****03_L and 03_S are distinct regions of interest (ROIs) from the same donor tissue section that were defined prior to MERSCOPE aquisition and subsequently combined into a single sample for downstream analysis.**

HuP-3 (HuP67A) = 568,761 cells

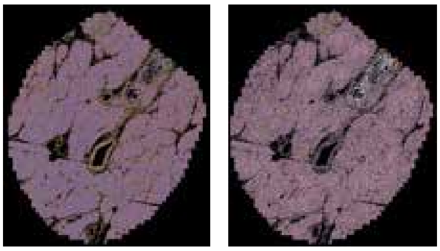

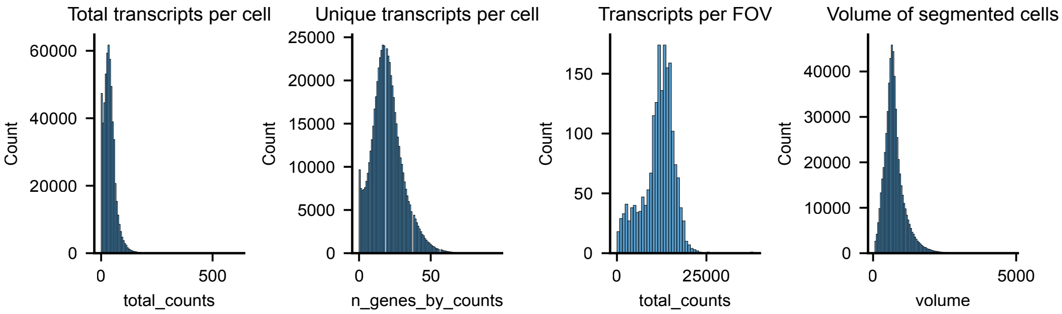


HuP-2 (HuP20A) = 670,736 cells

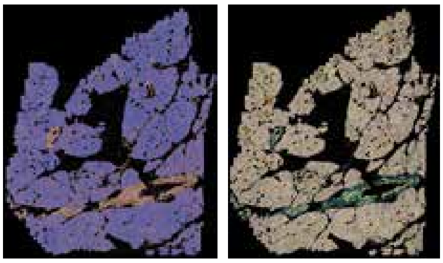

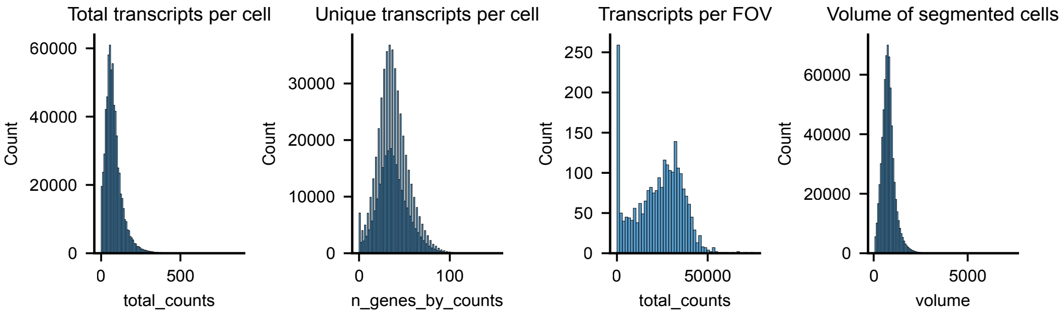

### Supplementary Figure 2

**B continued.**

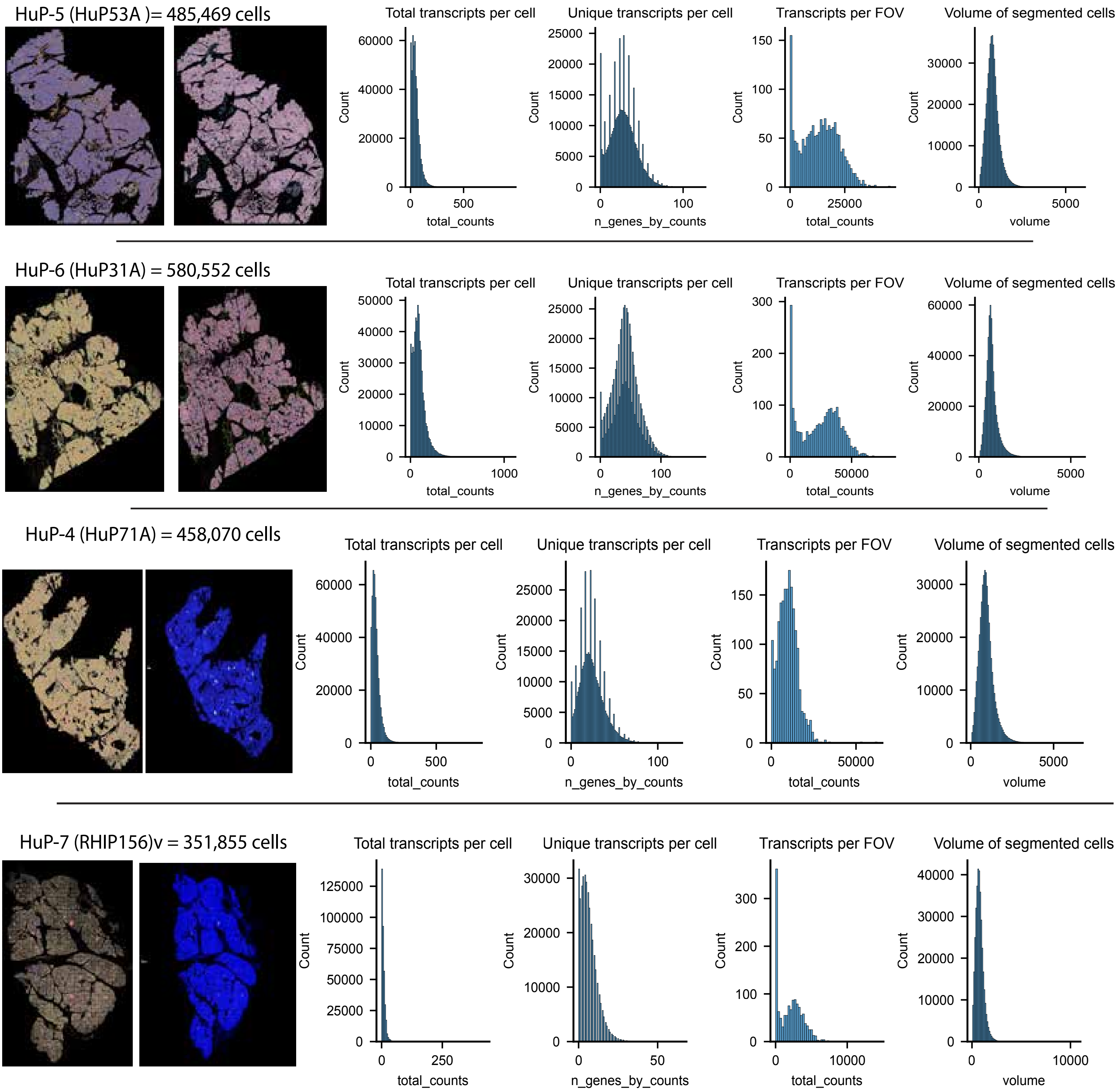


C

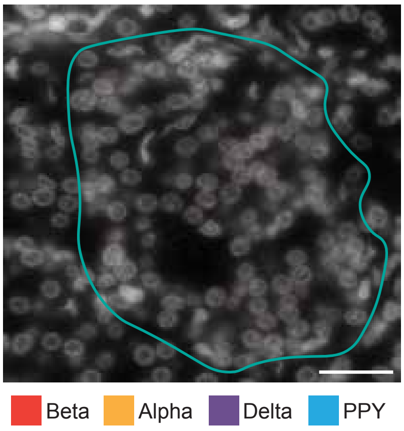


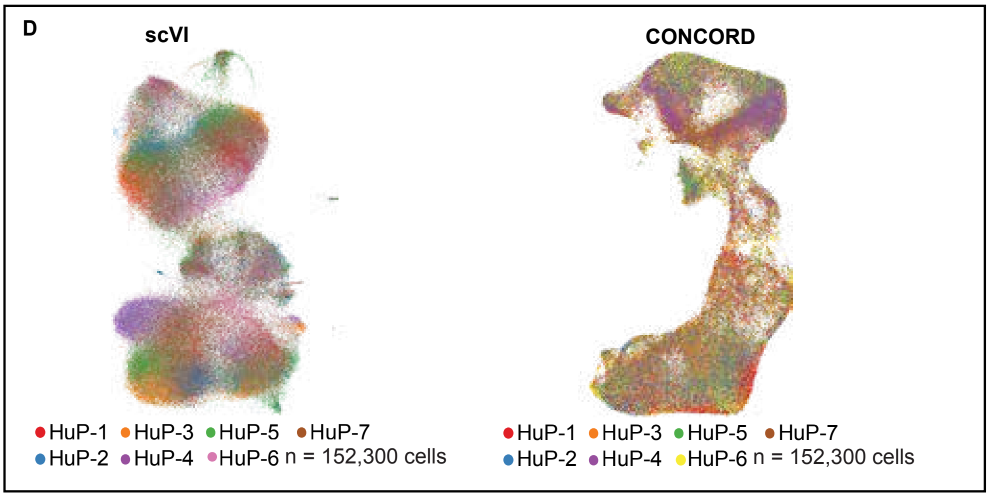

### Supplementary Figure 2

E

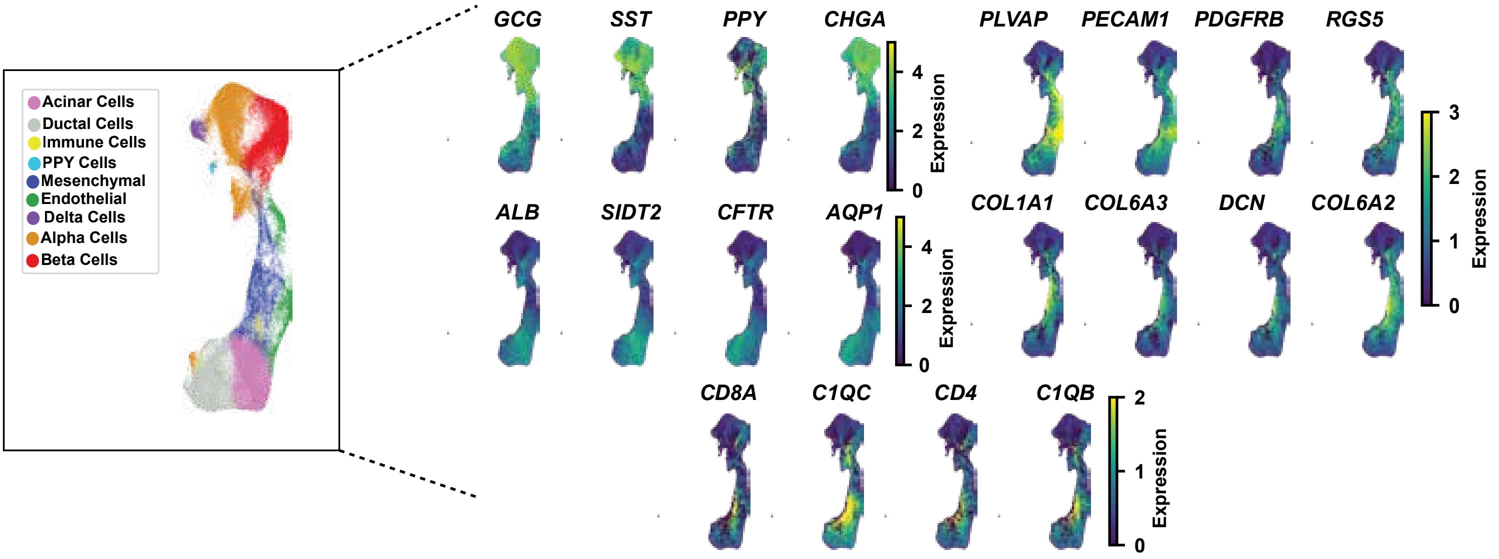


F

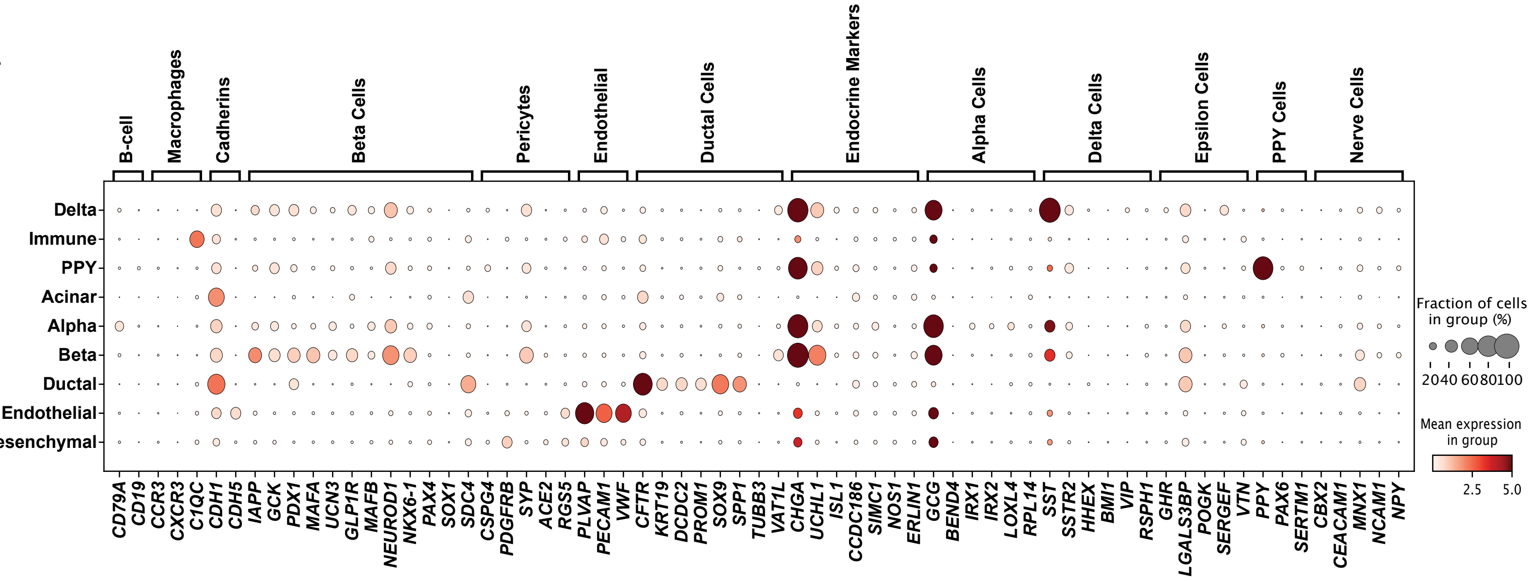


G

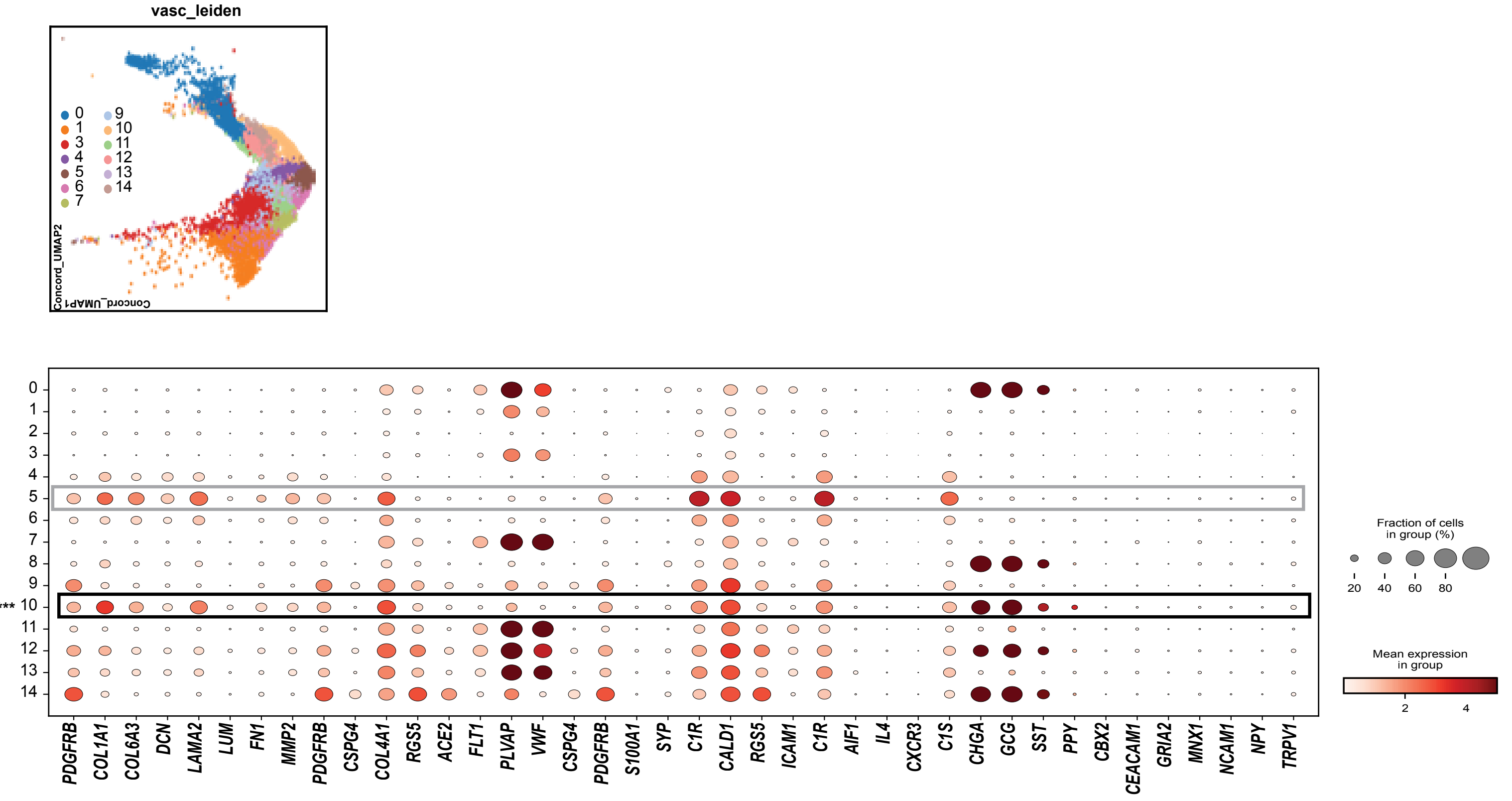

### Supplementary Figure 2

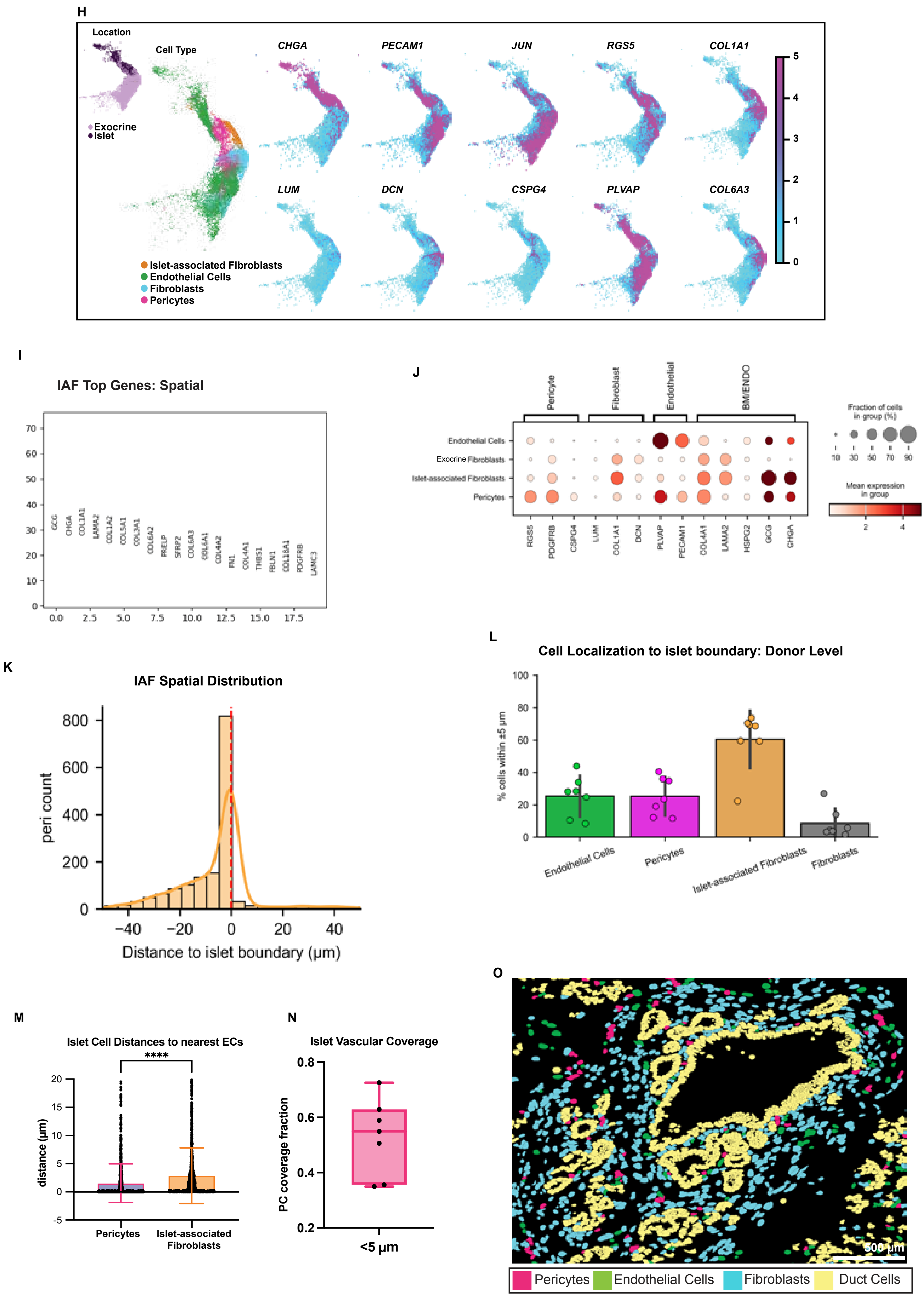

### Supplementary Figure 3

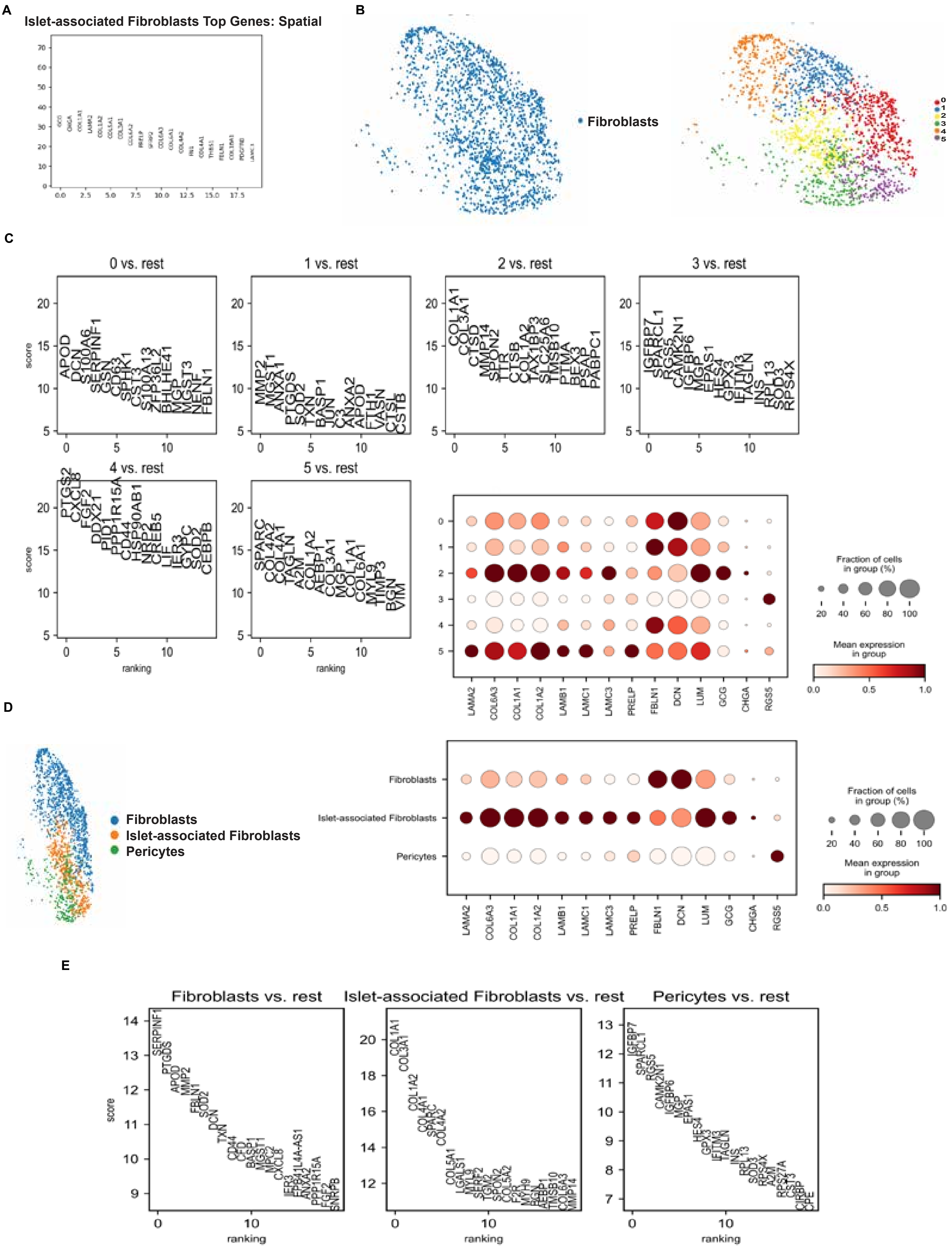

#### Supplementary Figure 3

**F**

**Craig-Schapiro Dataset**

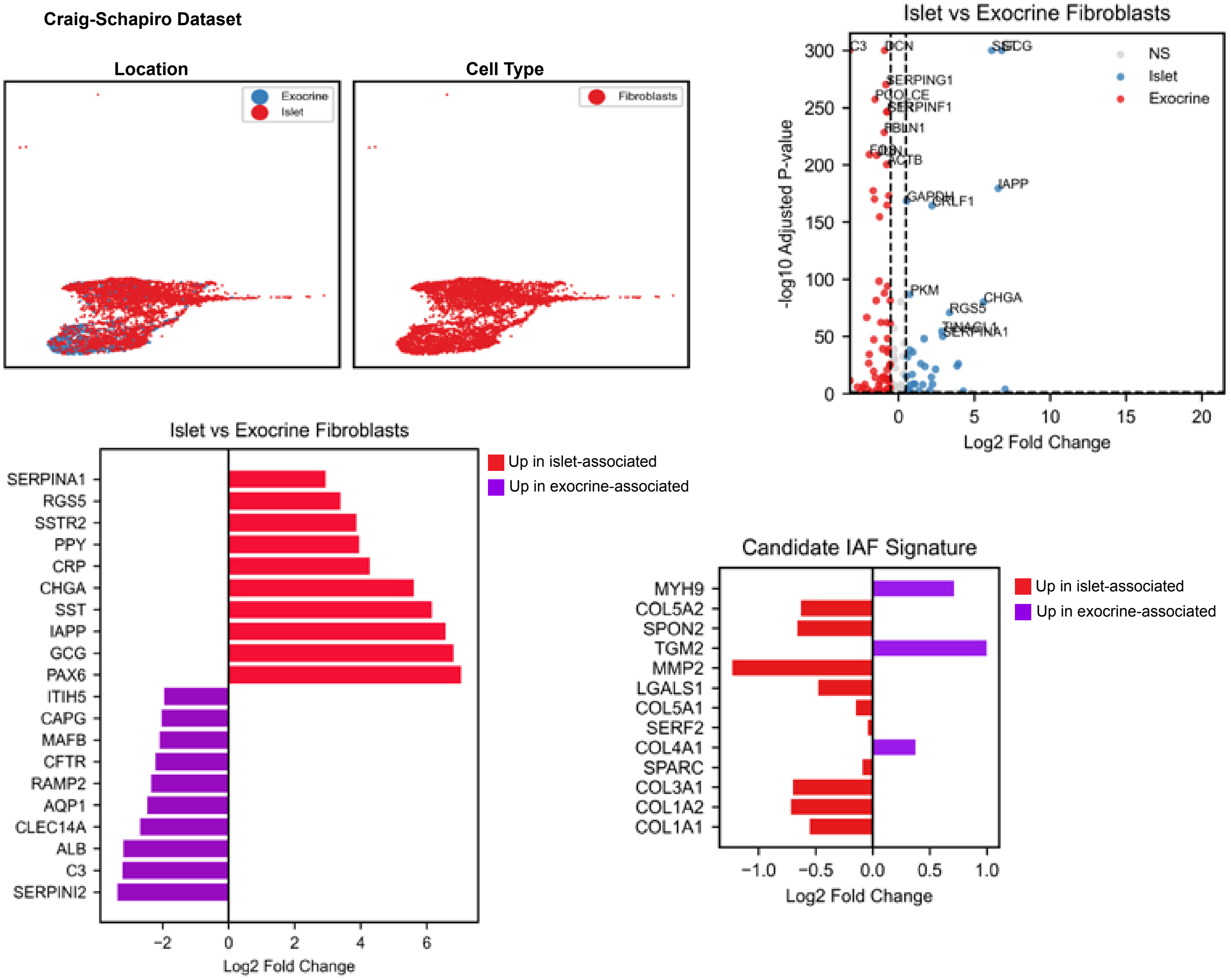


**G** **Craig-Schapiro Dataset**
**Islet-associated Fibroblasts**
**Pearson r = 0.608, p = 1.11e-31**
**Spearman = 0.664, p = 1.78e-39**

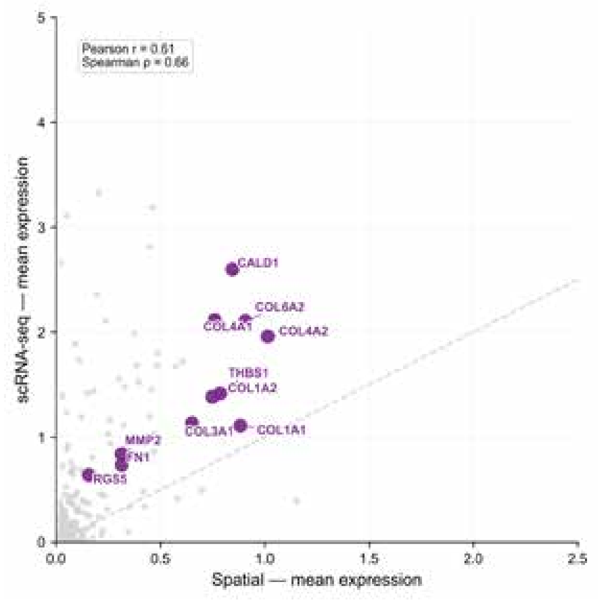

### Supplementary Figure 4

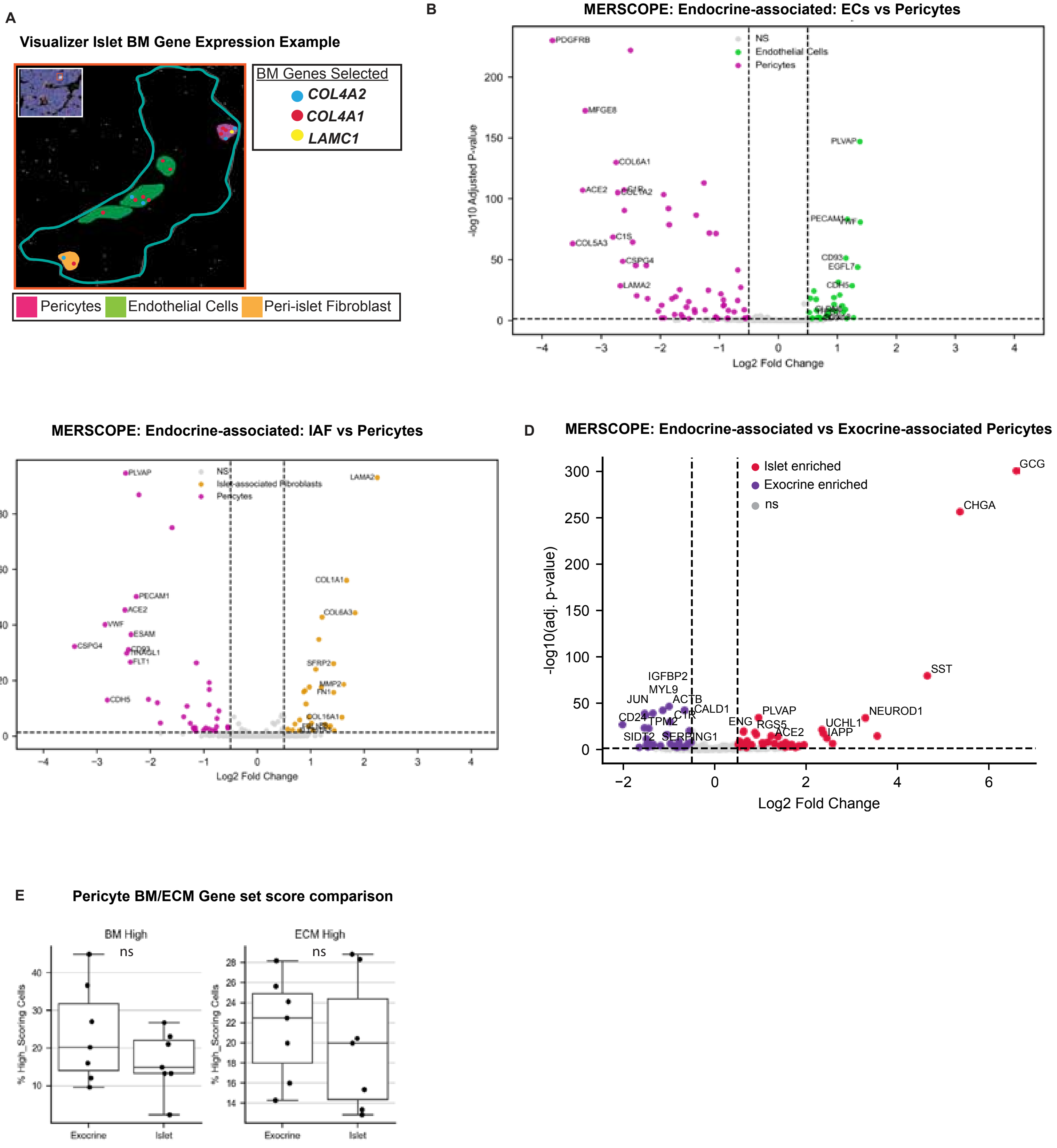

### Supplementary Figure 5

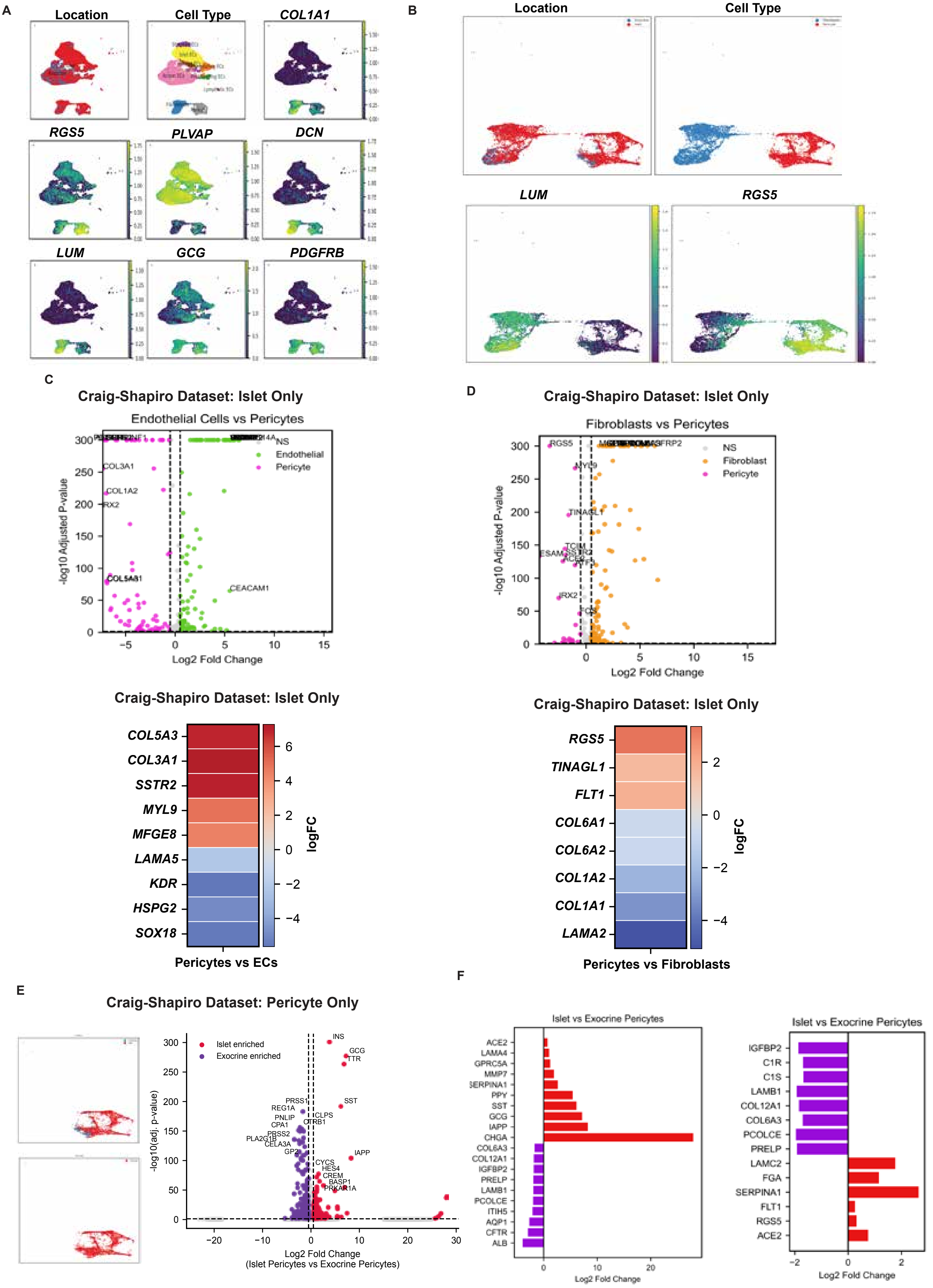

### Supplementary Figure 6

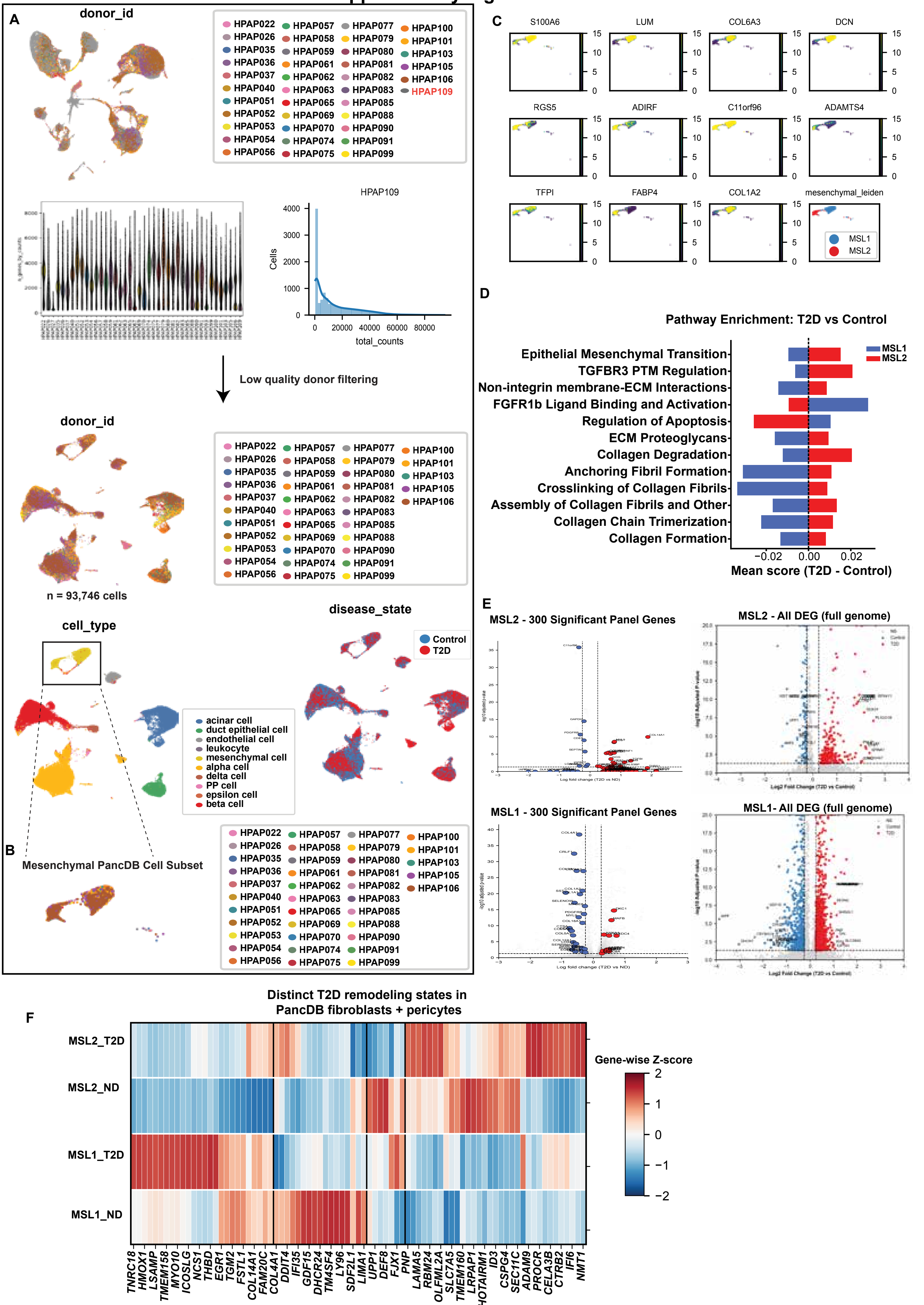

### Supplementary Figure 6

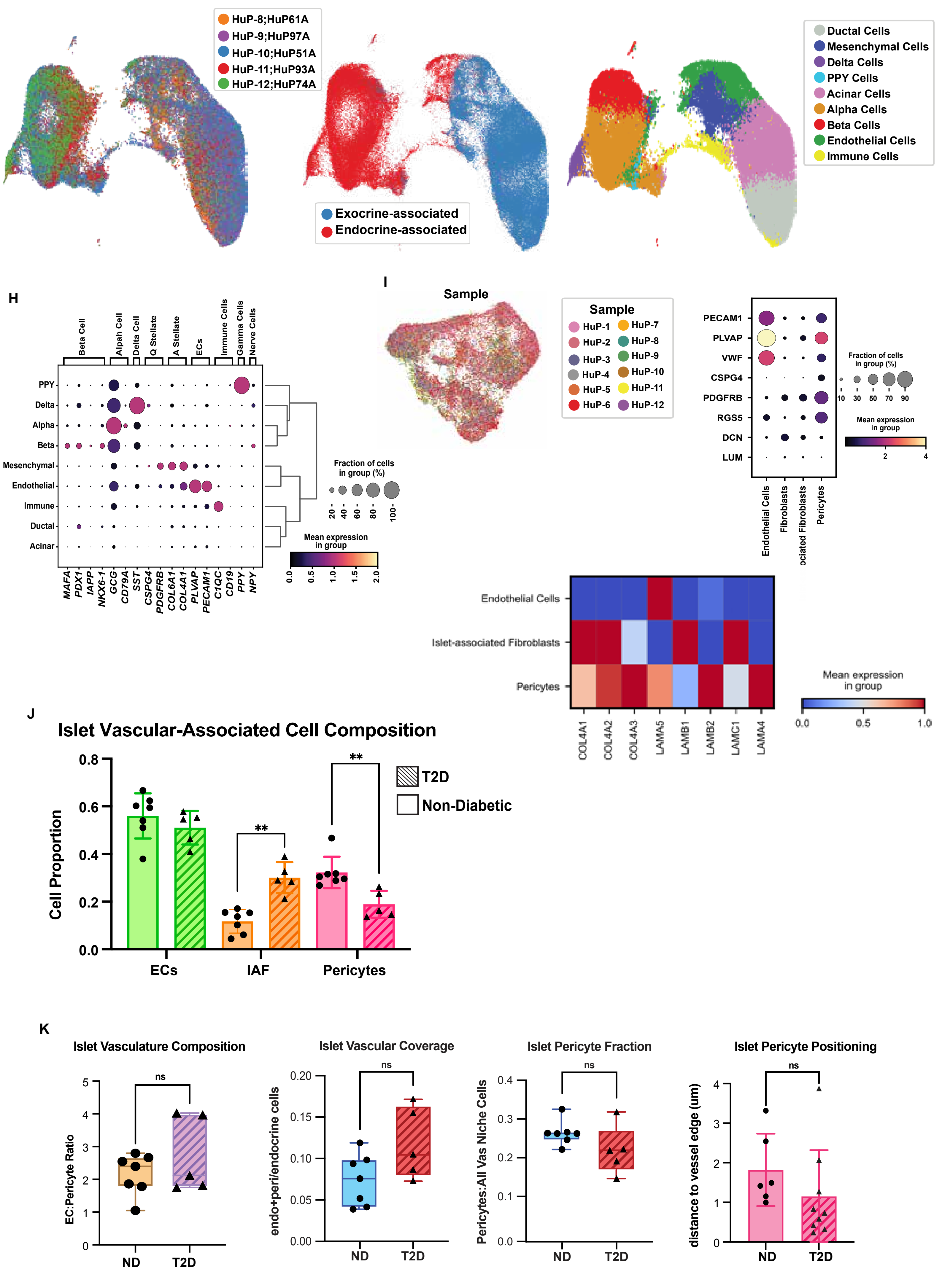

### Supplementary Figure 6

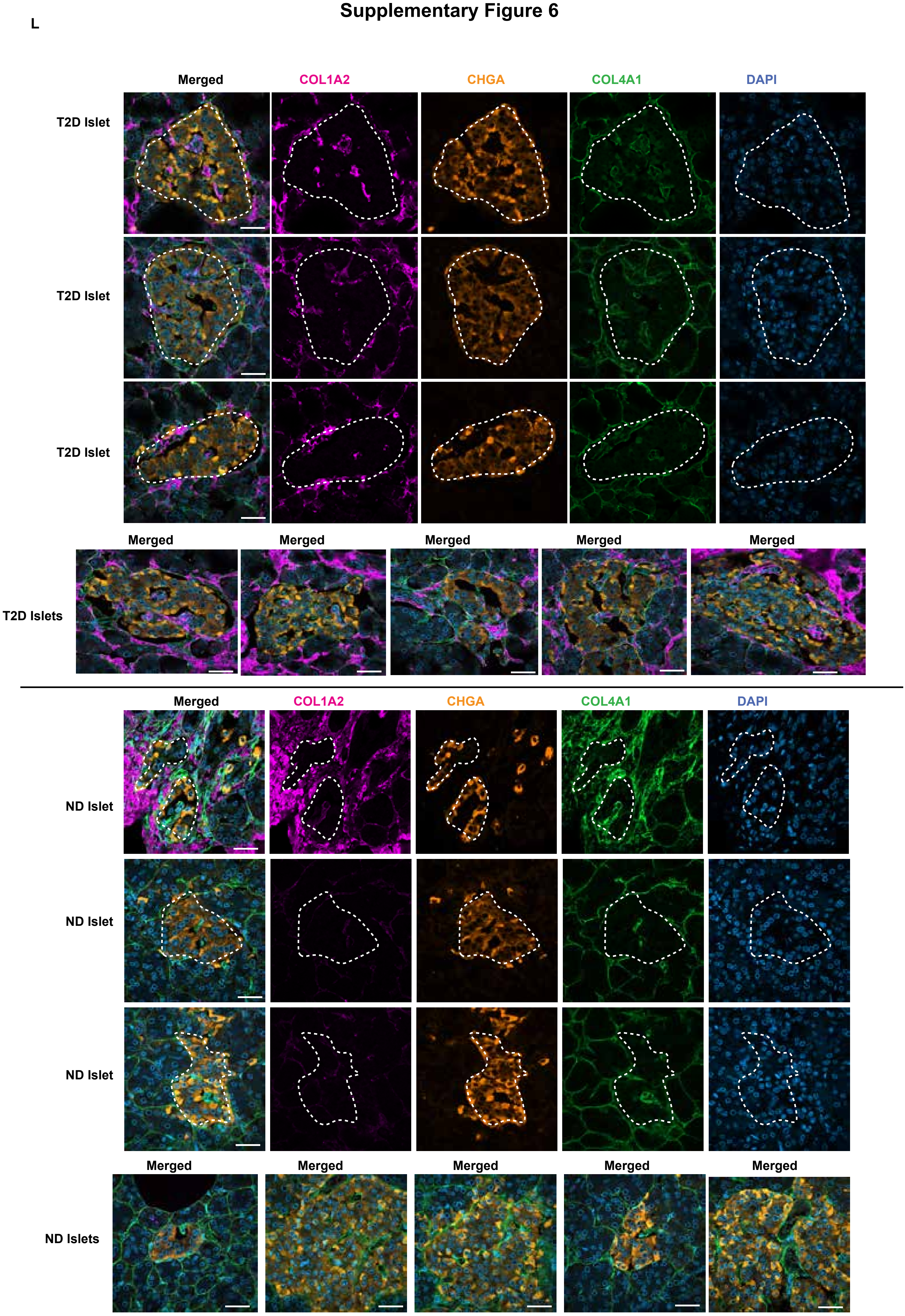
